# Postharvest properties of ultra-late maturing peach cultivars and their attributions to *Melting Flesh* (*M*) locus: Re-evaluation of *M* locus in association with flesh texture

**DOI:** 10.1101/2020.10.02.324582

**Authors:** Ryohei Nakano, Takashi Kawai, Yosuke Fukamatsu, Kagari Akita, Sakine Watanabe, Takahiro Asano, Daisuke Takata, Mamoru Sato, Fumio Fukuda, Koichiro Ushijima

**Affiliations:** Experimental Farm of Graduate School of Agriculture, Kyoto University, Kizugawa, Kyoto, Japan; Graduate School of Environmental and Life Science, Okayama University, Okayama, Japan; Faculty of Food and Agricultural Sciences, Fukushima University, Fukushima, Japan

**Keywords:** fruit, softening, ethylene, *Prunus persica*, melting flesh locus, endoPG, postharvest

## Abstract

The postharvest properties of two ultra-late maturing peach cultivars, ‘Tobihaku’ (TH) and ‘Daijumitsuto’ (DJ), were investigated. Fruit were harvested at commercial maturity and held at 25°C. TH exhibited the characteristics of normal melting flesh (MF) peach, including rapid fruit softening associated with an increase in endogenous ethylene production In contrast, DJ did not soften at all during three-week experimental period even though substantial ethylene production was observed. Fruit of TH and DJ were treated with 5000 ppm of propylene, an ethylene analog, continuously for seven days. TH softened rapidly whereas DJ maintained high flesh firmness in spite of an increase in endogenous ethylene production, suggesting that DJ but not TH lacked the ability to be softened in response to endogenous and exogenous ethylene/propylene. DNA-seq analysis showed that tandem endo-polygalacturonase (*endoPG*) genes located at *melting flesh* (*M*) locus, *Pp-endoPGM* (*PGM*) and *Pp-endoPGF* (*PGF*), were deleted in DJ. The *endoPG* genes at *M* locus are known to control flesh texture of peach fruit, and it was suggested that the non-softening property of DJ is due to the lack of *endoPG* genes. On the other hand, TH possessed an unidentified *M* haplotype that is involved in determination of MF phenotype. Structural identification of the unknown *M* haplotype, designated as *M^0^*, through comparison with previously reported *M* haplotypes revealed distinct differences between *PGM* on *M^0^* haplotype (*PGM-M^0^*) and *PGM* on other haplotypes (*PGM-M^1^*). Peach *M* haplotypes were classified into four main haplotypes: *M^0^* with *PGM-M^0^*; *M^1^* with both *PGM-M^1^* and *PGF*; *M^2^* with *PGM-M^1^*; and *M^3^* lacking both *PGM* and *PGF*. Re-evaluation of *M* locus in association with MF/non-melting flesh (NMF) phenotypes in more than 400 accessions by using whole genome shotgun sequencing data on database and/or by PCR genotyping demonstrated that *M^0^* haplotype was the common haplotype in MF accessions, and *M^0^* and *M^1^* haplotypes were dominant over *M^2^* and *M^3^* haplotypes and co-dominantly determined the MF trait. It was also assumed on the basis of structural comparison of *M* haplotypes among *Prunus* species that the ancestral haplotype of *M^0^* diverged from those of the other haplotypes before the speciation of *Prunus persica*.

## Introduction

Fruit firmness is an important quality that influences consumer preference, damage during distribution, and shelf life. Studies associated with the decrease in fruit firmness after harvest have been conducted with an eye toward reducing distribution loss and prolonging shelf life and thus, supplying high-quality fruit to consumers (Carrasco-Valenzuela et al., 2019; Fernandez I Marti et al., 2018; Liu et al., 2018; Moggia et al., 2017; Nimmakayala et al., 2016; Tucker et al., 2017). Fruit can be classified as climacteric or non-climacteric depending on their respiration and ethylene production patterns during ripening (Biale and Young, 1981). In climacteric fruit, ethylene is acknowledged to play an important role in controlling ripening- and senescence-related phenomena including fruit softening, due to the fact that massive ethylene production commences at the onset of ripening; exogenously applied ethylene and/or ethylene analog, propylene, induces ripening and senescence; ethylene inhibitors retard the progress of fruit ripening and senescence; and mutants and transgenic lines defective in ethylene production ability exhibit suppressed fruit ripening, especially softening (Gapper et al., 2013; Minas et al., 2015; Tucker et al., 2017).

Peach (*Prunus persica* (L.) Batsch) is generally known to belong to the climacteric type and to exhibit dramatic increases in respiration and ethylene production during ripening (Tonutti et al., 1991). In melting flesh (MF) peaches, the increased ethylene stimulates fruit softening principally through cell wall modification (Brummell et al., 2004; Hayama et al., 2006, 2008; Liu et al., 2018). MF peaches are highly perishable, softening rapidly after harvest. The increasing interest in improving peach shelf life has sparked investigations and resulted in findings of peach strains with long shelf lives. Those studies have demonstrated phenotypic variability associated with fruit softening and identified the possible causal genes for peach shelf life, as described below.

Intensively studied peaches that have long shelf lives are the stony-hard (SH) and slow-ripening (SR) peaches (Haji et al., 2005; Brecht and Kader, 1984; Bassi and Monet, 2008). The SH is determined by *Hdhd* gene and fruit with SH flesh bear the *hdhd* genotype (Haji et al., 2005). SH peaches are characterized by the absence of ethylene production and high firmness during postharvest storage, which are caused by the reduced expression of ethylene biosynthesis related gene *PpACS1* encoding 1-aminocyclopropane-1-carboxylic acid synthase (Tatsuki et al., 2006). YUCCA flavin mono-oxygenase gene *PpYUC11*, which is involved in the auxin biosynthesis pathway, has been proposed as a candidate for causal gene for this phenotype (Pan et al., 2015; Tatsuki et al., 2018). SR peaches are known to show delayed maturation on the tree, thereby resulting in late harvest. In SR peaches harvested earlier than the optimum harvest date, flesh firmness decreased slowly (Brecht and Kader, 1984). Its genetic base was characterized and a deletion mutation in a gene encoding the NAC transcription factor was reported to be responsible for the SR phenotype (Eduardo et al., 2015; Meneses et al., 2016; Nuñez-Lillo et al., 2015).

Another peach strain that shows high flesh firmness during postharvest ripening is non-melting flesh (NMF) peaches (Fishman et al., 1993; Yoshioka et al., 2011). Whereas MF peaches soften dramatically and bear melting texture during the final stage of ripening called “melting phase”, NMF fruit appear to lack this “melting phase” of softening and remain relatively firm during ripening not only on the tree and but also after harvest (Fishman et al., 1993; Yoshioka et al., 2011). The MF/NMF phenotypes segregate as a single locus (*M*) that is linked tightly to the stone adhesion locus (Bailey and French 1949; Monet, 1989). MF is dominant over NMF and the recessive allele determines the NMF character (Bailey and French, 1949; Monet, 1989). In NMF peaches, solubilization of cell wall pectin and enzymatic activity and protein accumulation of endo-polygalacturonase (endoPG), a pectin hydrolase, are markedly reduced compared with MF peaches (Fishman et al., 1993; Lester et al., 1996; Pressey and Avants, 1978; Yoshioka et al., 2011). Studies aimed at demonstrating *endoPG*(*s*) as a candidate gene for *M* locus have shown suppressed or undetectable expression of *endoPG*(*s*) in NMF peaches and polymorphisms in *endoPG* genes coinciding with MF/NMF phenotypes (Callahan et al., 2004; Gu et al., 2016; Lester et al., 1994, 1996; Morgutti et al., 2006, 2017; Peace et al., 2005, 2007). *M* locus is located at 3.5 cM interval on the bottom of linkage group 4 of the peach map, the position within which a genomic region with clusters of *endoPG* genes exists (Cao et al., 2016; Gu et al., 2016). Two tandem *endoPG* genes in that region, *Pp-endoPGM (PGM*) and *Pp-endoPGF (PGF*), corresponding to sequences *Prupe.4G2622OO* in v2.0 of peach genome (*ppaOO6857m* in v1.0) and *Prupe.4G2619OO (ppaOO6839m* in v1.0), respectively, were found to be responsible for determining the MF/NMF phenotypes (Gu et al., 2016). Gu et al. (2016) proposed a scenario where *M* locus has three allelic copy number variants of *endoPG* genes designated by H_1_ (possessing *PGF* and *PGM*), H_2_ (only *PGM*), and H_3_ (*null*). Accessions harboring either H_1_ and/or H_2_ haplotype (H_1_H_1_, H_1_H_2_, H_1_H_3_, H_2_H_2_, H_2_H_3_) exhibit MF phenotype whereas those harboring homozygous recessive H_3_ (H_3_H_3_) show NMF phenotype (Gu et al., 2016). It was also speculated that H_2_ is the ancestral haplotype whereas H_1_ and H_3_ haplotypes are two variants due to the duplication and deletion of *PGM*, respectively (Gu et al., 2016). However, research on different NMF peach germplasms suggested that mutations in *endoPG* gene(s) could be of more than one type arising from more than one source (Callahan et al., 2004; Lester et al., 1996; Morgutti et al., 2006, 2017; Peace et al., 2005) and some NMF accessions seemed to be incompatible with the model proposed by Gu et al. (2016). Much more comprehensive evaluation of *M* locus in association with flesh textural traits is required. Pursuing *M* locus evolution with much broader genetic resources covering *Prunus* species is also necessary for precise judgement.

Peach is known to have high diversity with regard to not only flesh texture but also fruit maturation date (Elsadr et al., 2019). In Japan, MF peaches reaching maturation stage in early July through September are mainly produced. Recently, because of the increasing demand for fresh peach in late autumn, ultra-late maturing cultivars whose optimum harvest dates are October and November are gathering the attention of growers. However, the postharvest properties of some of these rare cultivars have not yet been characterized.

In this study, first, two postharvest properties, namely, ethylene production and fruit softening, of two extremely late harvest cultivars, ‘Tobihaku’ (TH) and ‘Daijumitsuto’ (DJ), were investigated. It was revealed that TH showed normal MF peach ripening properties, whereas DJ possessed unique properties in that the fruit did not soften at all in spite of significant endogenous ethylene production and exogenous propylene treatment. Second, DNA-seq analysis of these cultivars demonstrated that DJ was a homozygote of an allele lacking *PGM* and *PGF* at *M locus*, whereas TH possessed a previously structurally unidentified haplotype that contained one *endoPG*. Third, accessing database sequences at *M* locus in more than 400 peach accessions and *Prunus* species indicated that the newly identified haplotype was an important allele that distributed widely within MF accessions, determined MF phenotype, and seemed to have been diverse from the other haplotypes before the speciation of *P. persica*. The scenario is discussed in which not three but four allelic variants at *M* locus are associated with the flesh texture and the newly identified haplotype is one of the two dominant determinants of MF texture.

## Materials and Methods

### Plant Materials

Ultra-late maturing peach (*Prunus persica* (L.) Batsch) cultivars ‘Tohbihaku’ (TH) and ‘Daijumitsutoh’ (DJ), whose genetic backgrounds are unknown, were examined. TH fruit were harvested on November 7, 2018, the commercial harvest date, from trees grown in a commercial orchard in Okayama Prefecture located in southwestern Japan. DJ fruit were harvested on October 12, 2018, the commercial harvest date, from trees grown in the Research Farm of the Faculty of Agriculture, Okayama University. As regards DJ, fruit from a different production area, namely, Fukushima Prefecture, which is located in northeastern Japan, were harvested on October 22, 2018, the commercial harvest date, and used to investigate the effects of growing conditions and harvest maturity. Climate conditions in the peach production areas, which were obtained from the website of the Japan Meteorological Agency (http://www.jma.go.jp/jma/index.html), are listed in Table S1. Regardless of cultivar or growing area, fruit were grown under suitable climate conditions for peach production by skilled growers using conventional growing techniques in Japan, including fruit thinning and bagging by the end of June. The fruit were harvested on the basis of skilled growers’ visual evaluation using de-greening of fruit ground color as harvest index. After harvest, fruit were ripened at 25°C for three weeks and ethylene production rate, flesh firmness, soluble solids content (SSC), and juice pH were measured on days 0, 7, 14, and 21 at 25°C. DJ fruit harvested from Fukushima Prefecture were packed carefully in a corrugated cardboard box and transported by vehicle at ambient temperature to Okayama University two days after harvest, where fruit were ripened at 25°C.

### Propylene Treatment

Some fruit were treated with propylene, an ethylene analog, continuously for seven days at 25°C, as described previously (Hiwasa et al., 2004; McMurchie et al., 1972). Ethylene production rates and flesh firmness were measured on treatment days 0, 3, and 7.

### Measurement of Ethylene Production Rate, Flesh Firmness, SSC, and Juice pH

Ethylene production rate, flesh firmness, SSC, and juice pH were measured as described previously (Kawai et al., 2018; Nakano et al., 2018). For the measurement of ethylene production rate, individual fruit were incubated in a 1.3-liter plastic container at room temperature for 30 min. Headspace gas withdrawn from the container was injected into a gas chromatograph (GC8 CMPF; Shimadzu, Kyoto, Japan) equipped with a flame ionization detector (set at 200°C) and an activated alumina column (φ 4 mm × 1 m) set at 80°C. For flesh firmness, the cheek parts of each fruit were cut and peeled, and flesh penetration force was measured using a rheometer (FUDOH RTC Rheometer; RHEOTECH, Tokyo, Japan) with a 3-mm-diameter cylindrical plunger and expressed in Newton per plunger area (N/mm^2^). The relationships between the flesh penetration force measured by this system and fruit maturity indexes are listed in Table S2. SSC and juice pH in the cheek parts of each fruit were measured with a refractometer (PR-1; Atago, Tokyo, Japan) and a pH meter (B-712; HORIBA, Kyoto, Japan), respectively.

### Statistical Analysis

Three to four fruit were used as biological replicates at each measurement point. Data with significant differences of the means were evaluated using Tukey’s multiple comparison test.

### Mapping WGS Data and Variant Calling

Genomic DNA was isolated from the leaves of DJ, TH, and ‘Benihakutoh’ (BH) by Nucleon PhytoPure (Cytiva). DNA-seq analysis was performed by Novogen and 9G data of PE150 reads, corresponding to approx. 30 times coverage, were obtained. We further searched SRA (Sequence Read Archive) database to obtain whole genome shotgun sequencing (WGS) data for doubled haploid ‘Lovell’ (dhLL), Dr. Davis (DD), and ‘Big Top’ (BT). Illumina WGS reads were mapped to peach reference genome (ver. 2.0) by CLC Genomics Workbench or minimap2 (Li, 2018). SNP, indel, and structural variants were called by CLC Genomics Workbench.

### De Novo Assembly and Structural Comparison

DNA-seq analysis of BH was further performed by Macrogen to obtain 46G data of PE150 reads, which correspond to approx. 170 times coverage of *P. persica* genome. Illumina reads were assembled by ABySS 2.0 (Jackman et al., 2017). Contigs encompassing *M* locus were detected by Blastn analysis using *PG1, PG2, PGM/F, NADH*, and *F-box* genes, which were located in the *M* locus region, as query. *M^0^* and *M^1^* haplotype sequences (see Results) were compared by nucmer (Kurtz et al., 2004) and their relationship was drawn by Circos (http://circos.ca). We generated new reference sequence set “PpREF20+M0”, in which the *M^0^* haplotype sequence was added to *P. persica* reference genome. Illumina short reads of DJ, TH, BH, dhLL, DD, and BT were mapped to PpREF20+M0 by minimap2. Coverage was analyzed from BAM file by samtools and drawn by Circos.

### Genotyping by PCR

Genomic DNA was extracted from leaves of peach accessions. Genotyping by PCR was conducted with six sets of primers described in Table S3. PCR was performed with BIOTAQ DNA Polymerase (Bioline, UK) using the following program: 30-35 cycles at 95°C for 20 s, annealing for 15 s, and extension at 72°C an initial denaturation at 95°C for 3 min, and a final extension at 72°C for 7 min. Annealing temperature and extension time are described under ‘PCR conditions’ in Table S3. PGM/F products were separated on a 15% acrylamide gel and others were separated on an agarose gel. PCR products were stained with UltraPower DNA Safedye (Gellex International Co., Ltd., Japan).

## Results

### Fruit Ripening Characteristics of Two Ultra-Late Maturing Cultivars

Postharvest changes in ethylene production rate, flesh firmness, SSC, and juice pH were investigated in two ultra-late maturing peach cultivars, TH and DJ. In TH, as much as 22 nl·g^-1^·h^-1^ of ethylene was produced at harvest and the amount increased gradually during storage at 25°C (Fig. 1A). Flesh firmness was 0.7 N/mm^2^ at harvest but decreased dramatically to less than 0.2 N/mm^2^ by day 7 at 25°C (Fig. 1D). Thereafter, the decrease became slight until the last day of experiment (day 21). SSC and juice pH did not change remarkably during storage (Fig. S1). In DJ harvested from the Research Farm of Okayama University on October 12, ethylene production rate was almost negligible at harvest. Thereafter, ethylene production rate increased and peaked on day 7, reaching more than 10 nl·g^-1^·h^-1^ (Fig. 1B). Flesh firmness was around 0.85 N/mm^2^ at harvest (Fig. 1E). In spite of considerable ethylene production, flesh firmness showed no significant decrease during storage at 25°C and was almost unchanged until the last day of experiment (day 21). SSC slightly increased during storage, reaching a peak of 18 °Brix on day 14, and juice pH was maintained at around pH 4.0 during storage (Fig. S1).

**Figure 1.**
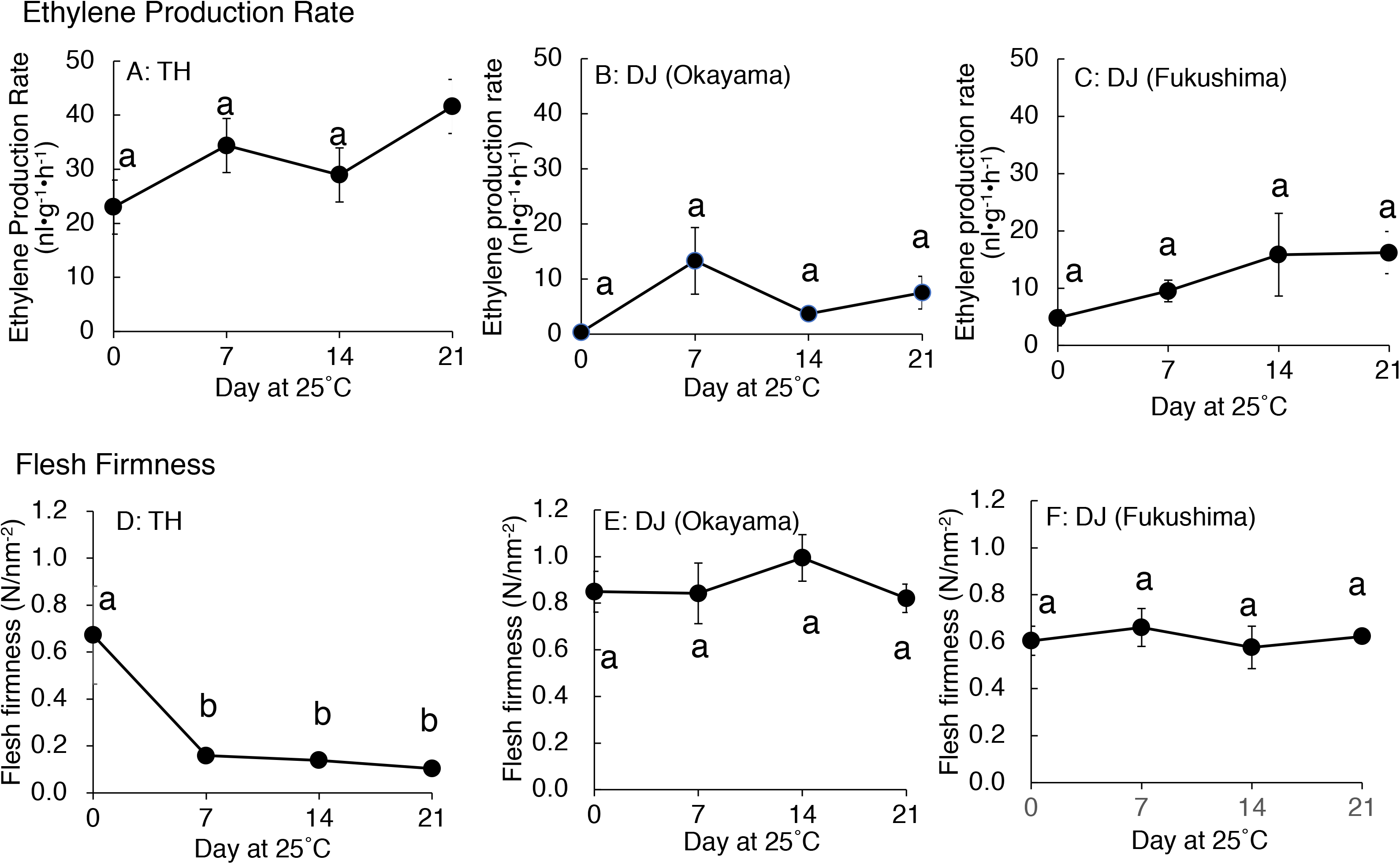
Postharvest changes in (A, B, C) ethylene production rate and (D, E, F) flesh firmness in ‘Tohbihaku’ (TH) and ‘Daijumitsutoh’ (DJ) fruit grown in Okayama Prefecture and DJ grown in Fukushima Prefecture Japan. In (A) and (D), TH were harvested on November 7 from a commercial orchard in Okayama Prefecture, Japan and held at 25°C for 21 days. In (B) and (E), DJ from Okayama were harvested on October 12 from the Research Farm of Okayama University, Japan and held at 25°C for 21 days. In (C) and (F), DJ from Fukushima were harvested on October 22 from a commercial orchard in Fukushima Prefecture, Japan, followed by two-day transport at ambient temperature to Okayama University, where fruit were held at 25°C for 21 days. Fruit were harvested at commercial maturity. Each point represents the mean value of three fruits and vertical bars indicate ± SE (n = 3 to 4). Different letters show significant difference by Tukey’s test at 5% level.

To confirm that the unique characteristics of DJ were not due to climatic effects and/or misestimated harvest maturity, DJ fruit harvested from a different production area were investigated. In the case of DJ harvested from a commercial orchard in Fukushima Prefecture on October 22, fruit delivered to Okayama University two days after harvest had an ethylene production rate of as high as 5.2 nl·g^-1^·h^-1^ and flesh firmness of 0.6 N/mm^2^ (Figs. 1C, F). Ethylene production rate increased during storage at 25°C, reaching 16 nl·g^-1^·h^-1^ on day 21. On the other hand, flesh firmness did not change significantly during storage; flesh firmness on day 21 was almost the same as that at harvest. SSC and juice pH did not change remarkably during storage (Fig. S1). Figure 2 shows DJ fruit on day 14. The external appearance showed no significant deterioration; however, the longitudinal sections showed internal browning and slight flesh breakdown around the stones.

**Figure 2.**
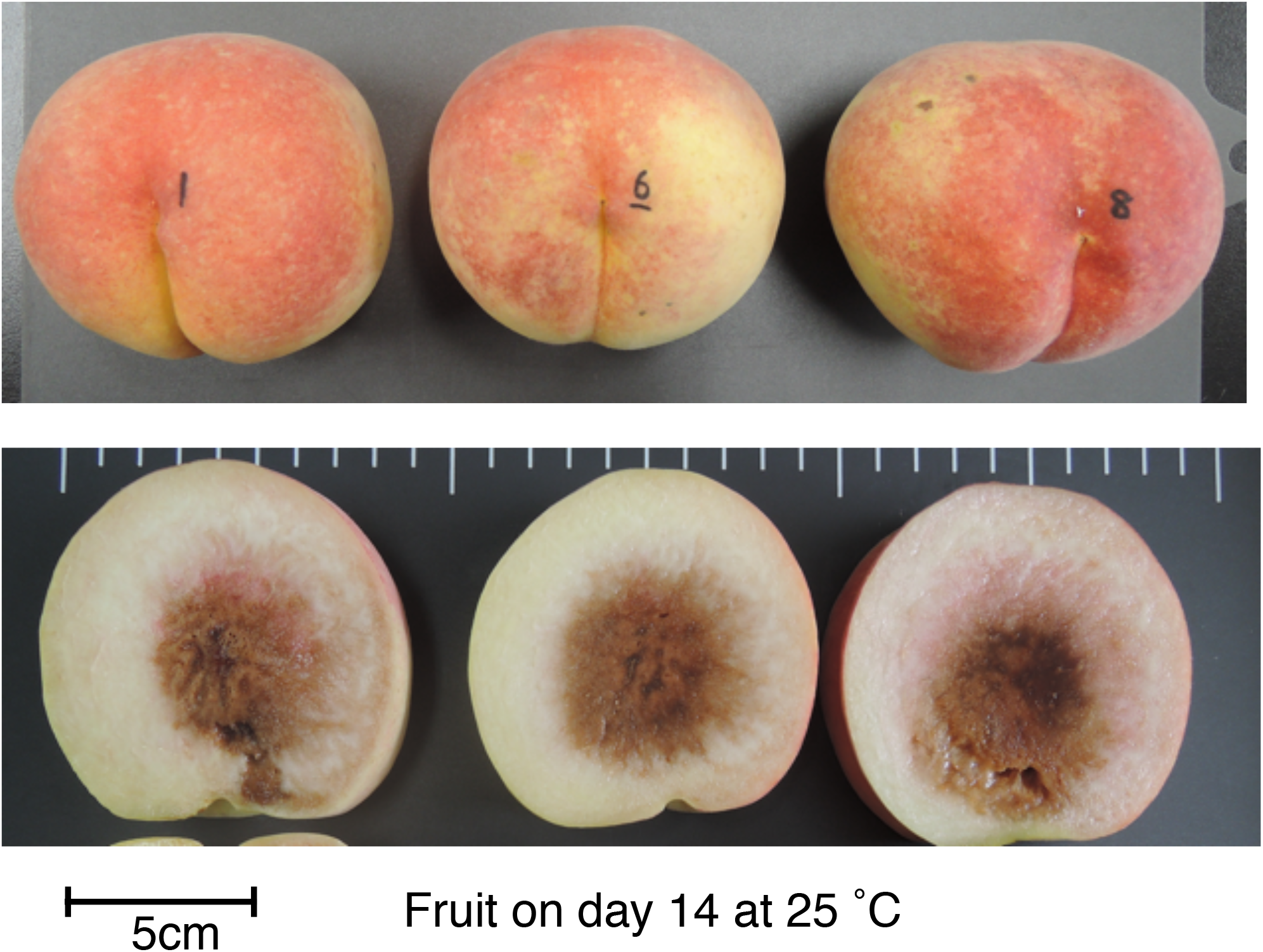
DJ on day 14 at 25°C. Fruit were harvested from a commercial orchard in Fukushima Prefecture and held at 25°C for 21 days, as shown in Figure 1. Appearance (top) and longitudinal section (bottom) of each fruit are shown. Bar indicates 5 cm.

### Different Responses to Exogenous Propylene Treatment of TH and DJ

Although ethylene production rate during storage was higher in TH than DJ, generally speaking, the amount of ethylene produced in DJ is sufficient to induce physiological effects on fruit ripening including flesh softening. In order to confirm that the non-softening characteristic of DJ is not due to the low ethylene production rate of this cultivar, TH and DJ were treated with 5,000 ppm of propylene, an ethylene analog, continuously for seven days. Ethylene production rates of TH and DJ were increased rapidly by the propylene treatment, reaching 45 and 22 nl·g^-1^·h^-1^ on day 3 of treatment, respectively (Fig. S2). Flesh firmness of TH decreased rapidly whereas that of DJ decreased slightly, and high firmness was maintained in DJ even after seven days of continuous propylene treatment (Fig. 3). These results suggest that regardless of the differences in the ethylene production rate, DJ lacked the ability to be softened in response to ethylene.

**Figure 3.**
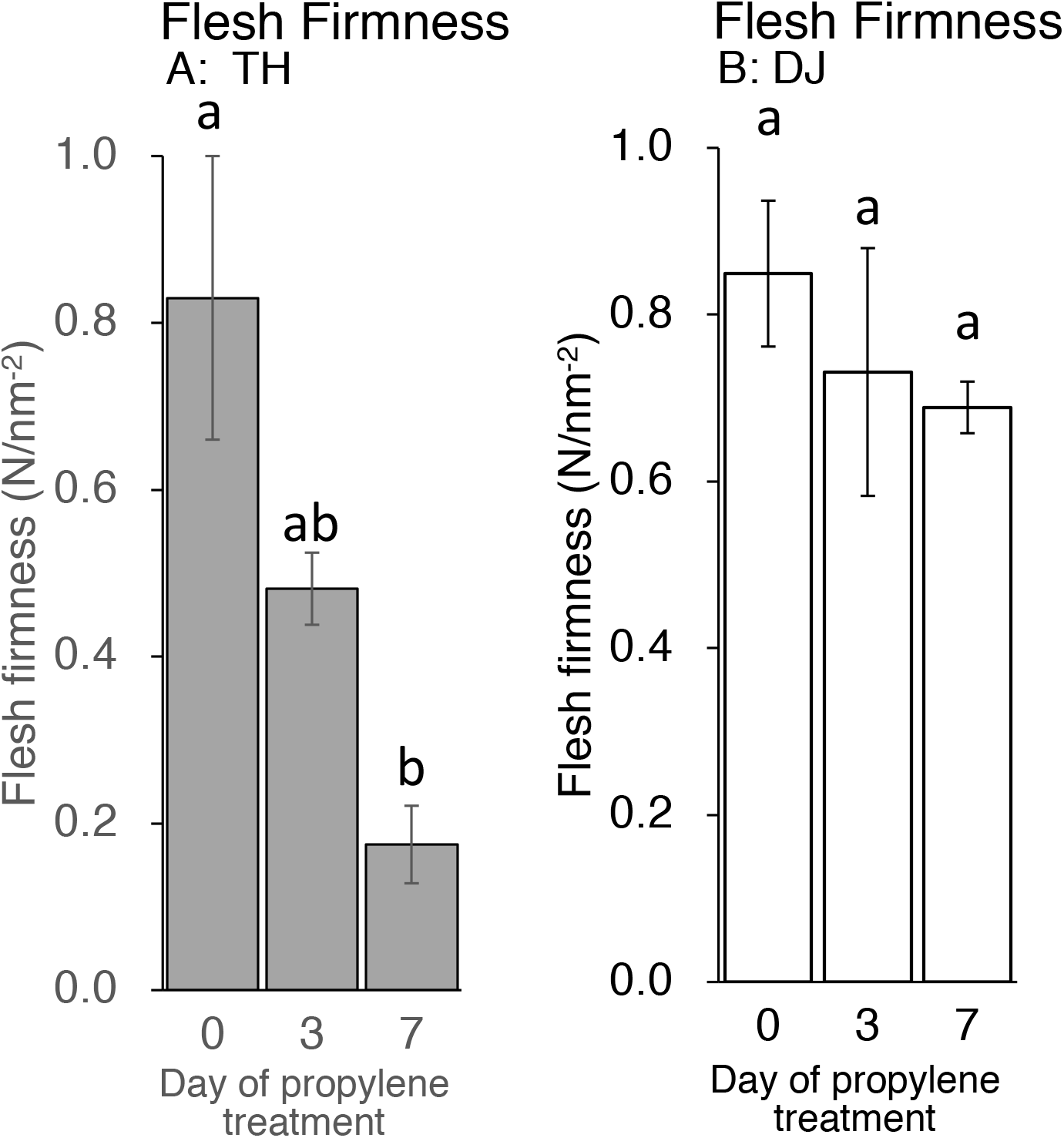
Effect of propylene treatment on postharvest fruit softening in (A) TH and (B) DJ. Harvested fruit were treated with 5,000 ppm of propylene continuously for seven days. Flesh firmness was measured on days 0, 3, and 7. For DJ, fruit harvested on October 12 from the Research Farm of the Faculty of Agriculture, Okayama University were used. Each point represents the mean value of three fruits and vertical bars indicate ± SE (n = 3 to 4). Different letters show significant difference by Tukey’s test at 5% level.

### Structural Features of *M* locus in DJ and TH and an Unidentified Haplotype in TH

*M* locus is involved in the regulation of peach flesh texture. Four polygalacturonase genes, *PG1, PG2, PGM*, and *PGF*; three *NADHs*; and one *F-box* gene were annotated in the *M* locus region of peach reference genome (Fig. 4A). Gu et al. (2016) walked the chromosome of three accessions by using PCR based on sequence information of reference genome, and proposed three haplotypes, H_1_, H_2_, and H_3_. The sequence of H_1_ haplotype was identical to reference genome, in which tandem duplication of *endoPG* genes, *PGM* and *PGF*, was observed. H_2_ haplotype possessed only *PGM* and lacked *PGF*. The 70 kbp region containing *PG2, PGM, PGF*, and *NADHs* was deleted in H_3_ haplotype. Gu et al. (2016) hypothesized that H_1_ and H_2_ haplotypes were dominant over H_3_ haplotype, *PGM* conferred the melting texture, and *PGF*, which was present only in H_1_ haplotype, was involved in freestone phenotype. The lack of both *PGM* and *PGF* in H_3_ homozygote could result in the non-melting texture. Further investigations are required to confirm the function of *PGM* in H_2_ and H_1_ haplotypes and the roles of *PGM* and *PGF* in fruit trait.

**Figure 4.**
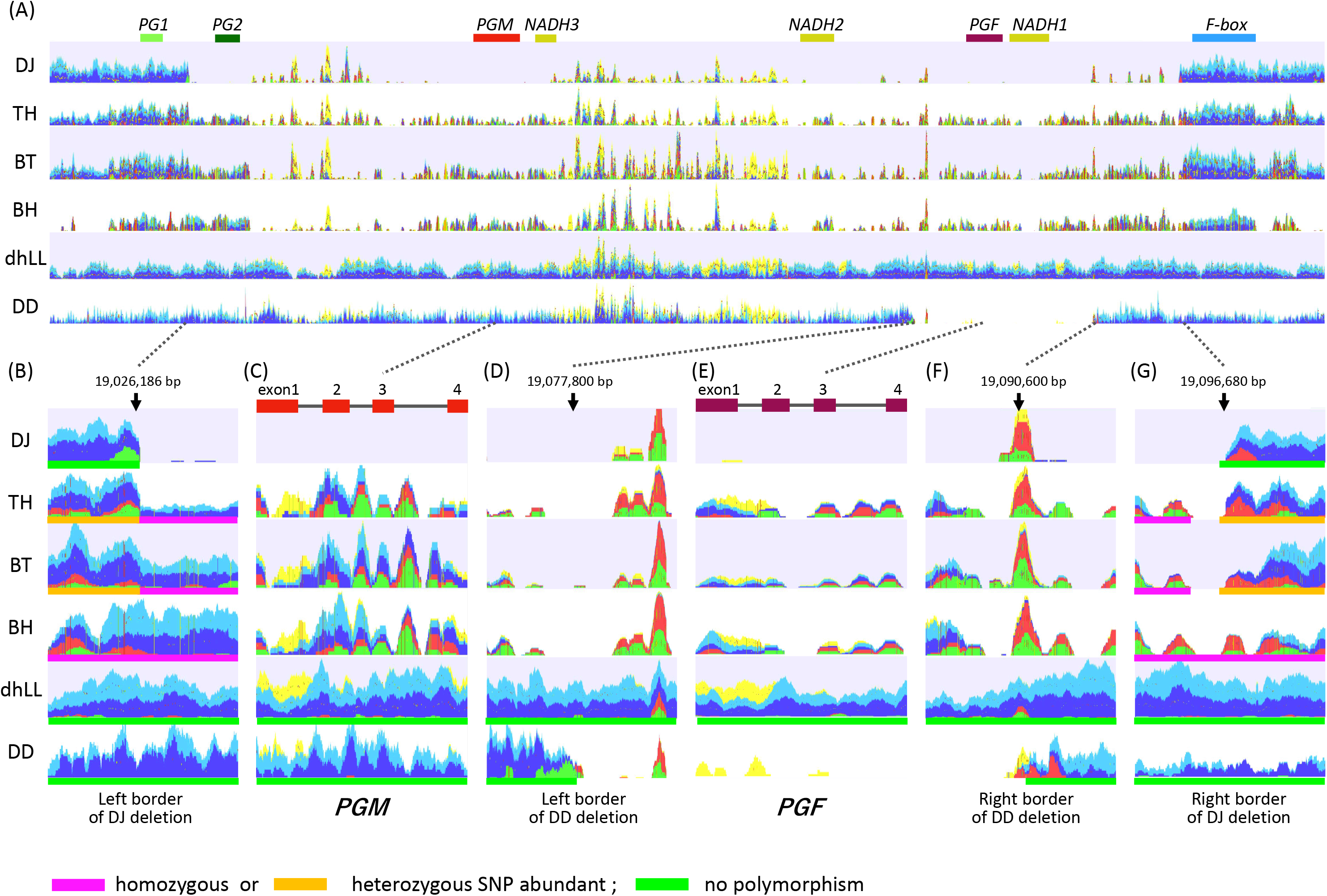
Mapping DNA-seq reads to *M^1^* haplotype. Illumina short reads of DJ, TH, BH, and dhLL were mapped to reference genome of *P. persica* (ver 2.0) by CLC Genomics Workbench. Properly paired reads that are in the correct orientation and distance are shown in blue and light blue. Green and red reads are broken pairs. Non-specific matches are shown in yellow. (A) Overview of mapping graph in the *M* locus region. *PG1* to *F-box* above graph are genes named according to Gu et al. (2016). (B) and (G) show the region around the junction of M3 deletion. (D) and (F) are those of M2 deletion. (C) and (E) are *PGM* and *PGF*, respectively.

WGS of DJ and TH by Illumina was performed to investigate the *M* locus region (19.0 to 19.1Mbp of chromosome 4) (Fig. 4), and the mapping patterns were compared with those of three accessions with different flesh textures: dhLL of MF accession, DD of NMF accession, and BT of MF accession, which is categorized in slow softening peaches (Morgutti et al., 2017; Ghiani et al., 2011; Bassi and Monet, 2008). dhLL, an H_1_ homozygote, was used for peach genome sequencing and the reads were mapped uniformly to *M* locus. The reads of DD were also mapped to the entire *M* locus region except the ~12 kbp region including *PGF* and *NADH1*. No or less reads were mapped to the ~12 kbp region, indicating that the region was deleted in DD. The position and length indicated that the DD deletion unexpectedly corresponded to the region deleted in H_2_ haplotype in Gu et al. (2016). The mapping patterns of DJ and TH were not the same as those of dhLL and DD, suggesting that DJ and TH possessed neither H_1_ nor H_2_ haplotype.

TH and BT showed similar mapping patterns and thus were expected to possess the same haplotypes (Fig. 4). The DJ mapping results indicated that the genotype of DJ was not identical to those of TH and BT, but the three accessions were expected to share one haplotype. Mapping the pair reads of DJ to peach reference genome revealed many broken pairs around 19,026,186 bp and 19,096,680 bp (Figs. 4A, B, E). Only a few reads were mapped between them, except for the many non-specific reads from other chromosomal regions that were mapped between *NADH3* and *NADH2*, the region named M1 insertion in this study (see below). Structural mutation analysis by CLC Genomics Workbench indicated that the region from 19,026,186 to 19,096,680 bp was deleted in DJ, suggesting that DJ is homozygote for the deleted haplotype identical to that reported as H_3_ by Gu et al. (2016). Many broken pairs were found at the same position as DJ when mapped with TH and BT pair reads, indicating that one of the two haplotypes was H_3_ (Figs. 4A, B, E). The other haplotype was presumed to be an unidentified novel haplotype that cannot be predicted directly by mapping to reference genome. In TH and BT, SNPs were heterozygous in the outer region of H_3_ deletion, whereas they were homologous in the inner region of H_3_ deletion (Figs. 4B, E, Table S4). All SNPs in this region were homozygous in BH, a Japanese MF accession. In contrast, no SNP was detected in either dhLL or DD. In the region of *PGM* and *PGF*, the coverage of TH, BT, and BH was low and many SNPs and broken pair reads were observed (Figs. 4C, D). Throughout *M* locus and its surrounding region, SNPs were hardly observed in dhLL, DD, and DJ, whereas many mutations were detected in TH, BT, and BH (Fig. 4, Table S4), suggesting the existence of an unidentified haplotype in TH, BT, and BH.

### Identification of *M^0^* Haplotype

As BH was presumed to be a homozygote of the unknown haplotype, de novo assembly with PE150 reads of BH was conducted and two contigs covering *M* locus were obtained. *PG1, PG2, PGM*, and *NADH* were located in one of the two contigs. The *F-box* gene was found in another contig. The haplotype composed of the two contigs was defined as *M^0^* according to the locus name. Haplotypes H_1_, H_2_, and H_3_ in Gu et al. (2016) were renamed *M^1^, M^2^*, and *M^3^*, respectively, based on haplotype features, to avoid confusion and define the haplotypes precisely.

When the genome structure of *M^0^* was compared with *M^1^* sequence, the region corresponding to *PGF* was not found in *M^0^* and only one *NADH* was located on *M^0^* (Fig. 5). The sequences were conserved among the two haplotypes but *M^1^* specific sequences were found in the region from *PG2* to *PGM* and the downstream region of *NADH3*. Many non-specific reads were mapped to the region spanning from *NADH3* to *NADH2* of*M^1^* haplotype, the region corresponding to M1 insertion described above (Figs. 4 and 5). The M1 insertion was assumed to be translocated from another chromosomal region and inserted into *NADH* to disrupt it, because both *NADH3* and *NADH2* were partial and their structures appeared to be generated from one *NADH* gene divided at the third intron by the M1 insertion (Figs. S3, S4 and NADH3/2 of Fig. S5).

**Figure 5.**
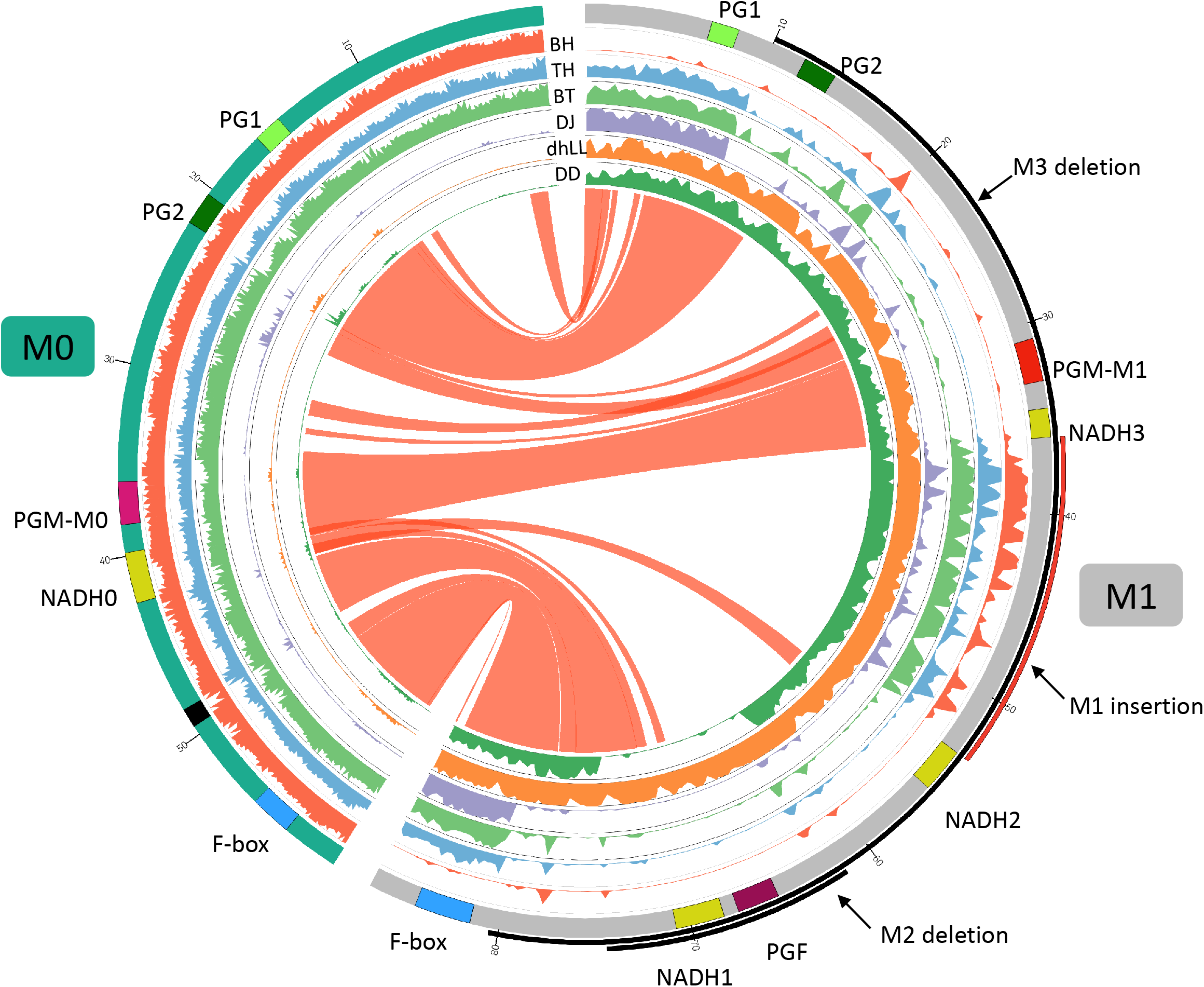
Comparison of *M^0^* and *M^1^* haplotypes. Inner six tracks are coverage graphs of DNA-seq reads of BH TH, BT, DJ, dhLL, and DD. Ribbons link similar regions between two haplotypes. *M^0^* haplotype consists of two contigs connected by N (black in the chromosome track). The nucleotide at 19,016,188 bp of Pp04 chromosome is positioned as +1 in *M^1^* haplotype. *PGM* of *M^0^* haplotype is similar to *PGM* of *M^1^* but not identical. We named those *PGMs PGM-M^0^* and *PGM-M^1^*, respectively. *NADH0* is *NADH* of *M^0^* haplotype and seems to be an intact gene as described in Figures S3, S4, and S5.

The CDS sequence of *PGM* of *M^0^* (*PGM-M^0^*) showed 99% similarity to those of *PGM* of M1 (*PGM-M^1^*) and *PGF* of *M^1^*. One amino acid substitution between *PGM-M^0^* and *PGM-M^1^* was found at residue 49, where Ser was substituted to Phe in *PGM-M^1^* (Fig. S6). The Ser at residue 49 of *PGM-M^0^* was conserved in other *Prunus* species. Amino acid sequence comparison between *PGM-M^0^* and *PGF* showed substitution of Ser for Thr at residue 269 in PGF, although it was not considered to have a significant effect on the function of the PG protein. The S49F substitution between f and fl alleles was reported by Peace et al. (2005) and Morgutti et al. (2017). However, because they did not perform resequencing analysis, it was not possible to understand the entire structure of the f haplotype and its origin. It was probable that f haplotype was the same as *M^0^* in this study. We further demonstrated that this *M^0^* haplotype is not a specific haplotype but a widely found haplotype in various peach accessions, and plays an important role in MF phenotype determination.

DNA-seq reads of BH, TH, DJ, dhLL, DD, and BT were mapped to the reference ‘PpREF20+M0’, in which the *M^0^* haplotype sequence was added to peach genome sequence (Fig. 5). Coverage graphs show that the reads of *M^0^, M^1^, M^2^*, and *M^3^* haplotypes could be classified clearly. The reads of BH (*M^0^M^0^*) were mapped only to *M^0^* haplotype and those of dhLL (*M^1^M^1^*) were also mapped only to *M^1^* haplotype. DJ (*M^3^M^3^*) and DD (*M^2^M^2^*) reads were mapped to *M^1^* haplotype except for the deleted region, as shown in Figure 4. The reads of TH (*M^0^M^3^*) and BT (*M^0^M^3^*) were mapped to both *M^0^* and *M^1^* haplotypes except for *M^3^* deletion.

### Structural Variety of *M* Haplotype Identified by WGS Mapping Patterns and PCR Genotyping

It was demonstrated that WGS data were useful for the precise genotyping of *M* locus. Many WGS data of peach accessions were registered in the SRA database and mapped to ‘PpREF20+M0’ to identify *M* genotype. We finally predicted *M* genotypes of 412 accessions from the WGS mapping pattern and/or by PCR genotyping. *M^0^* and *M^1^* haplotypes were the most popular in the peach accessions analyzed (Fig. 6, Table S5). On the other hand, the frequency of *M^2^* haplotype was very low. In addition to the four main haplotypes, *M^0^, M^1^, M^2^*, and *M^3^*, we found variant-type haplotypes and chimeric haplotypes. The former haplotypes exhibited deletion in the region different from those found in *M^2^* and *M^3^*, whereas the latter haplotypes appeared to be generated by the recombination between *M^0^* and *M^1^* (Fig. 6). In total, 11 haplotypes were structurally identified at *M* locus (Fig. 7). They were first classified into *M^0^* to *M^3^* on the basis of the existence of *PGM-M^0^, PGM-M^1^*, and *PGF*; the haplotype containing only *PGM-M^0^* is *M^0^*; the haplotype containing *PGM-M^1^* and *PGF* was *M^1^*; the haplotype containing only *PGM-M^1^* was *M^2^*; and the haplotype containing neither *PGM* nor *PGF* was *M^3^*. Furthermore, variants and recombinant types of haplotypes were identified from structural variations and such characters as b, c… or r1, r2… were added to their names, respectively.

**Figure 6.**
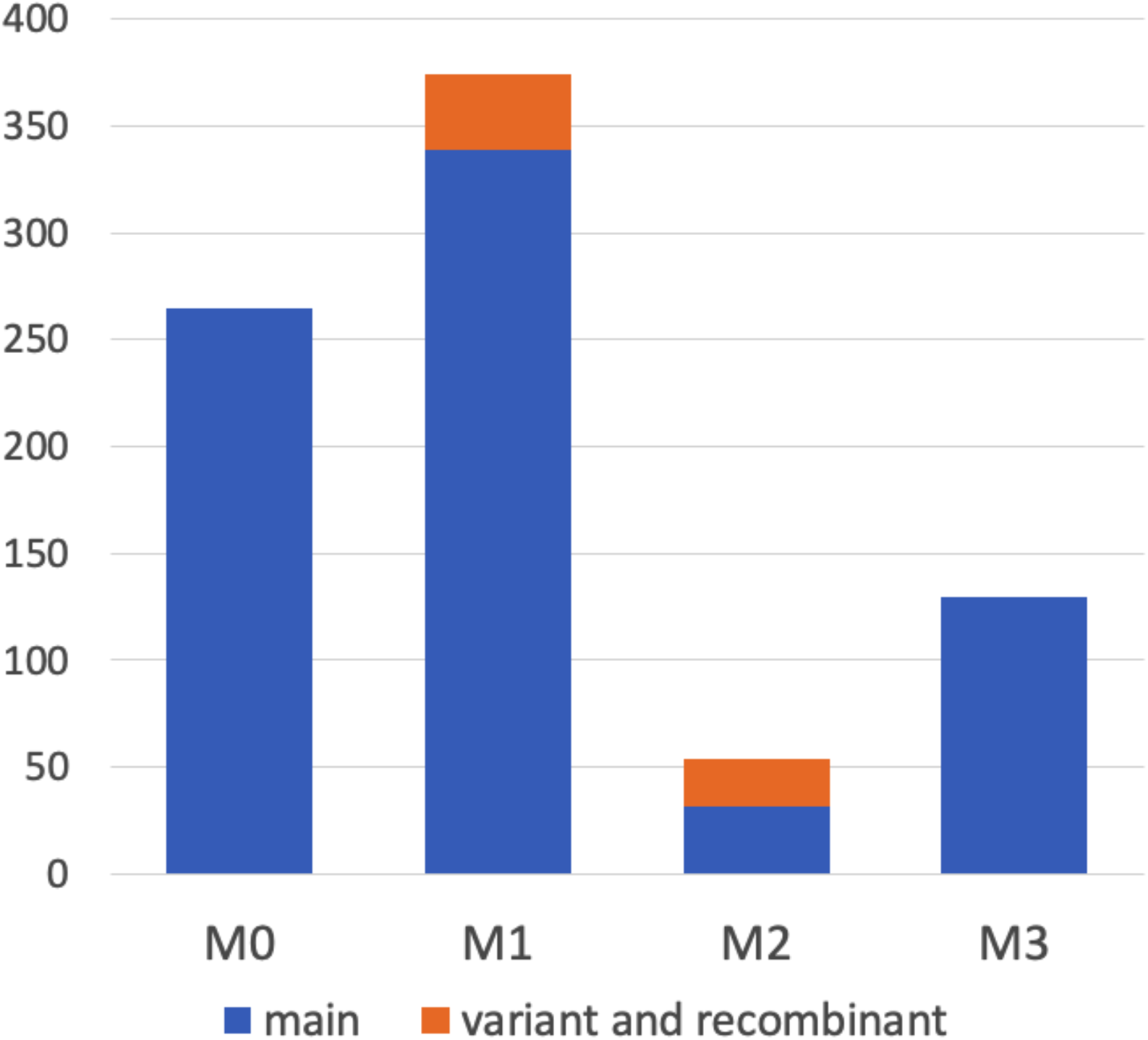
Haplotype frequency at *M* locus in peach accessions. We classified *M* haplotypes into 11 haplotypes (Fig. 7) in 412 peach accessions, on the basis of the coverage graph patterns and/or PCR genotyping (Figs. 8 and S9).

**Figure 7.**
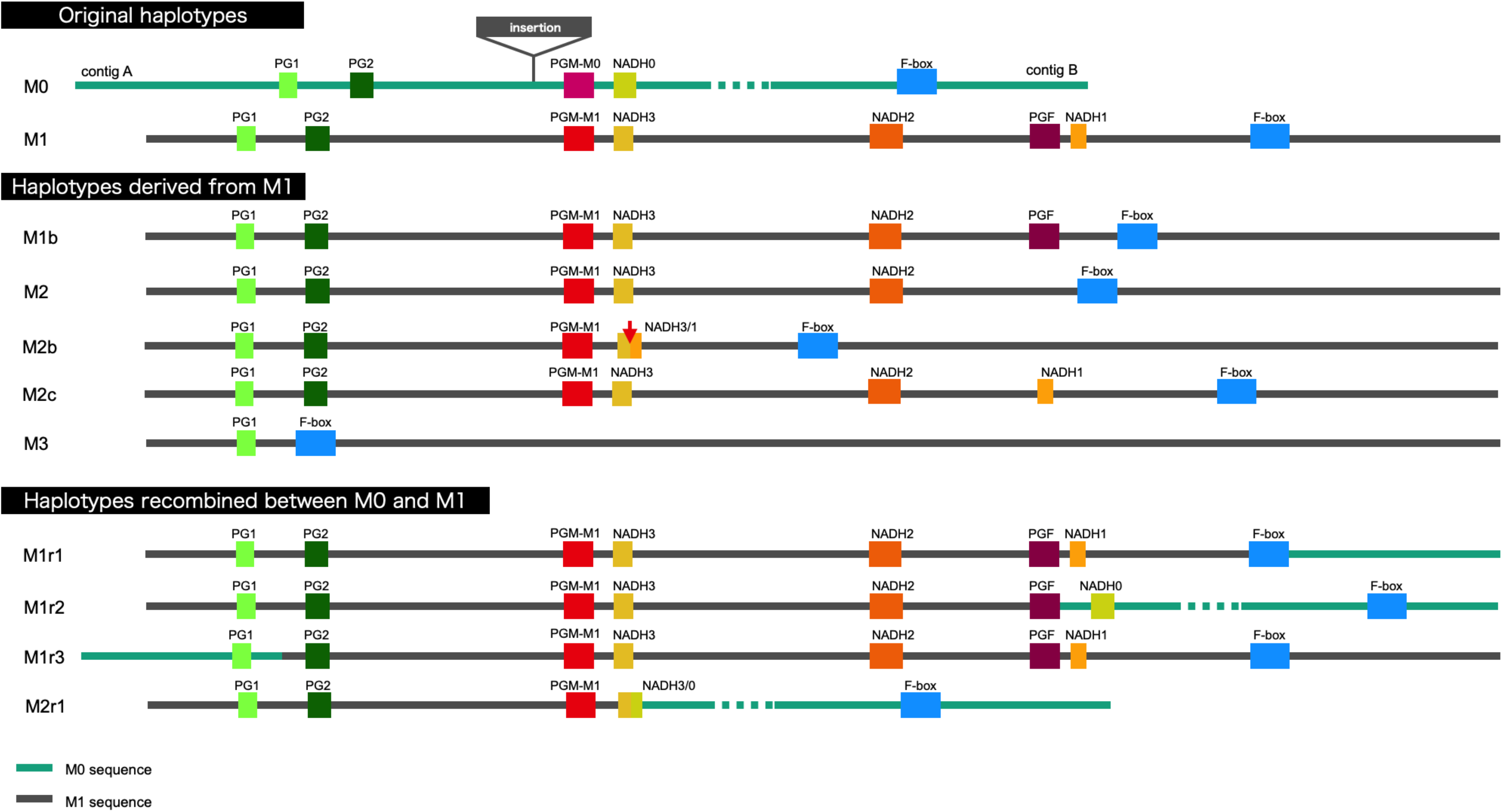
Schematic representation of *M* haplotypes of peach.

**Figure 8.**
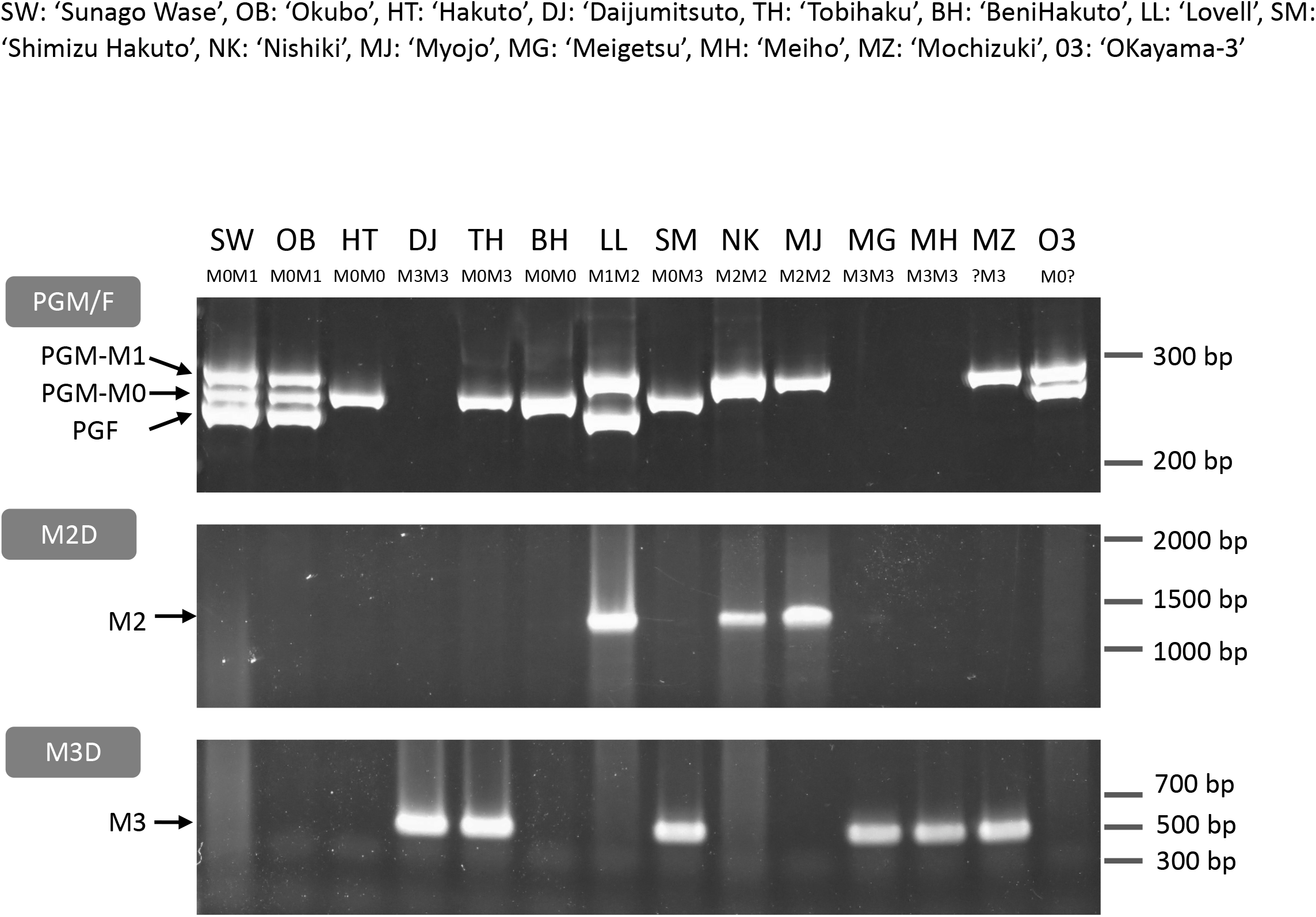
PCR genotyping of *M* locus. Three primer sets PGM/F, M2D, and M3D (Fig. S8, Table S3) were used to distinguish *M* haplotypes. PGM/F and M2D/M3D fragments were separated on an acrylamide gel and an agarose gel, respectively. PGM/F primer set amplified three fragments derived from *PGM-M^1^, PGM-M^0^*, and *PGF*, respectively. M2D and M3D detected *M^2^* and *M^3^* haplotypes, respectively. All genotypes under each cultivar name except MZ and O3 were identified from these fragment patterns (see Fig. S8). SW: ‘SunagoWase’, OB: ‘Okubo’, HT: ‘Hakuto’, DJ: ‘Daijumitsuto’, TH: ‘Tobihaku’, BH: ‘BeniHakuto’, LL: ‘Lovell’, SM: ‘ShimizuHakuto’, NK: ‘Nishiki’, MJ: ‘Myojo’, MG: ‘Meigetsu’, MH: ‘Meiho’, MZ: ‘Mochizuki’, O3: ‘Okayama-3’

Although no significant structural change in gene composition was detected, the sequence variation was found in the accessions with *M^0^* haplotype, when analyzed on the basis of the mapping patterns of DNA-seq reads of TH. The insertion, whose length was unknown, was found in the 1.8 kbp upstream region of *PGM-M^0^*, and SNPs were also detected in *PGM-M^0^*, one of which was located in CDS and led to a synonymous substitution (Fig. S7). It was expected that *M^0^* was also diversified. However, we regarded both types of *M^0^* as *M^0^* in this study because no changes in gene composition (structural feature as haplotype) and no amino acid substitutions were found.

### *M^0^* Haplotype was Widely Spread among MF Accessions

We designed primers for PCR genotyping on the basis of the sequence variations at the third intron of *PGM* and *PGF* and the *M* haplotype structural differences (Figs. 8 and S8, Table S3). The structural variations and the *PGM* and *PGF* compositions of *M* haplotypes identified in this study indicated that the four main haplotypes could be classified using three primer sets, PGM/F, M2D, and M3D. PGM/F primer set detected the indel at the third intron of *PGM* and *PGF*, and three different fragment sizes were amplified (Fig. 8). The upper fragment was derived from *PGM-M^1^*, the lower one was from *PGF*, and the middle one was from *PGM-M^0^*. When the middle fragment was amplified, the accession possessed *M^0^* haplotype. From *M^1^* haplotype, both the upper and the lower fragments should be amplified. When the lower fragment was not amplified and only the upper one was amplified, this meant that only *PGM-M^1^* was amplified, showing that the accession had *M^2^* haplotype. The genotyping results obtained with the PGM/F primer set could be confirmed by amplification with the M2D primer set that detects the deletion on *M^2^*. Because no fragment was amplified from *M^3^* with the PGM/F primer set, the M3D primer set, which amplifies the junction of *M^3^* deletion, should be useful to identify *M^3^* haplotype.

The three primer sets were used to genotype 14 accessions. The *M* genotypes of six accessions, ‘Okubo’ (OB), DJ, TH, BH, LL, and ‘Myojo’ (MJ), were predicted from the mapping patterns of WGS data, which were re-confirmed by PCR genotyping. In the other eight accessions, we found inconsistent amplification patterns in ‘Mochizuki’ (MZ) and ‘Okayama-3’ (O3). These two accessions were expected to have *M^2^* haplotype judging from the result that the PGM/F primer set amplified *PGM-M^1^* but not *PGF*. However, no amplification was observed in M2D. These results suggested that *M^2^* haplotype of MZ and O3 was a variant type of *M^2^*. Indeed, an additional primer set showed that they have M2b (Fig. S9).

Hakuto (HT), a progeny of ‘Chinese Cling’ (CC), was mainly used as germplasm for MF peach breeding in Japan (Fig. S10A). PCR genotyping showed that HT was a homozygote of *M^0^*, indicating the possibility that *M^0^* haplotype had been spread among Japanese peaches. All the major Japanese cultivars tested in this study were found to have *M^0^* haplotype (Fig. S10B). All cultivars except OB and ‘ShimizuHakuto’ (SM) were homozygotes of*M^0^*. SM was a typical MF cultivar in Japan, and its genotype was *MM^3^*, the same as that of TH (Fig. 8).

### *M^2^* and *M^3^* Could Confer NMF Phenotype

DJ was an *M^3^* homozygote and did not have *PGM* and *PGF*. Forty *M^3^* homozygotes were identified in this study. Flesh textural phenotypes of 16 out of 37 accessions were reported, and twelve accessions were reported as NMF except those whose flesh texture was only reported as supplemental data in Cao et al. (2016) (Table 1). This may support our conclusion that*M^3^* haplotype was likely involved in the determination of non-softening postharvest property in DJ. *M^2^* haplotype, in which only *PGM-M^1^* was located, also appeared to confer the NMF phenotype, unlike *M^0^* and *M^1^* haplotypes. In this study, we found 22 homozygotes for *M^2^* including variant types, 11 of which had fruit textural report(s), and all 11 were reported as NMF (Table 1). Only NJF16 accession had *M2M3* combination as well as phenotype report. NJF16 was reported to be NMF peach (Clark and Finn, 2010). MZ was *M^2b^M^3^* that had only *PGM-M^1^* as confirmed by PCR genotyping, and is a well-known Japanese NMF cultivar. All together, these findings suggest that *PGM-M^1^* may not be functional. On the other hand, *PGM-M^0^* and *PGF* seemed to confer the MF phenotype dominantly because flesh texture of *M^0^M^2^* (Tsukuba 86), *MM* (O3), *M^0^M^3^* (TH, SM, BT, and CC), *M^1^M^2^* (LL), and *M^1^M^3^* (‘Georgia Bell’ (GB)) was MF (Table 1). ‘Early Gold’ (EG) was identified as *M^1^M^1^* on the basis of the mapping patten of reads from the SRA database (Table S6), despite that the flesh texture was considered NMF (Yoshida, 1981). We suspected that the reads of EG registered in SRA were confused with those of the other accessions. This is because ‘Nishiki’ (K) (*M^2^M^2^*), an NMF accession, was the parent of EG and one of EG haplotypes was supposed to be *M^2^*.

**Table 1.**
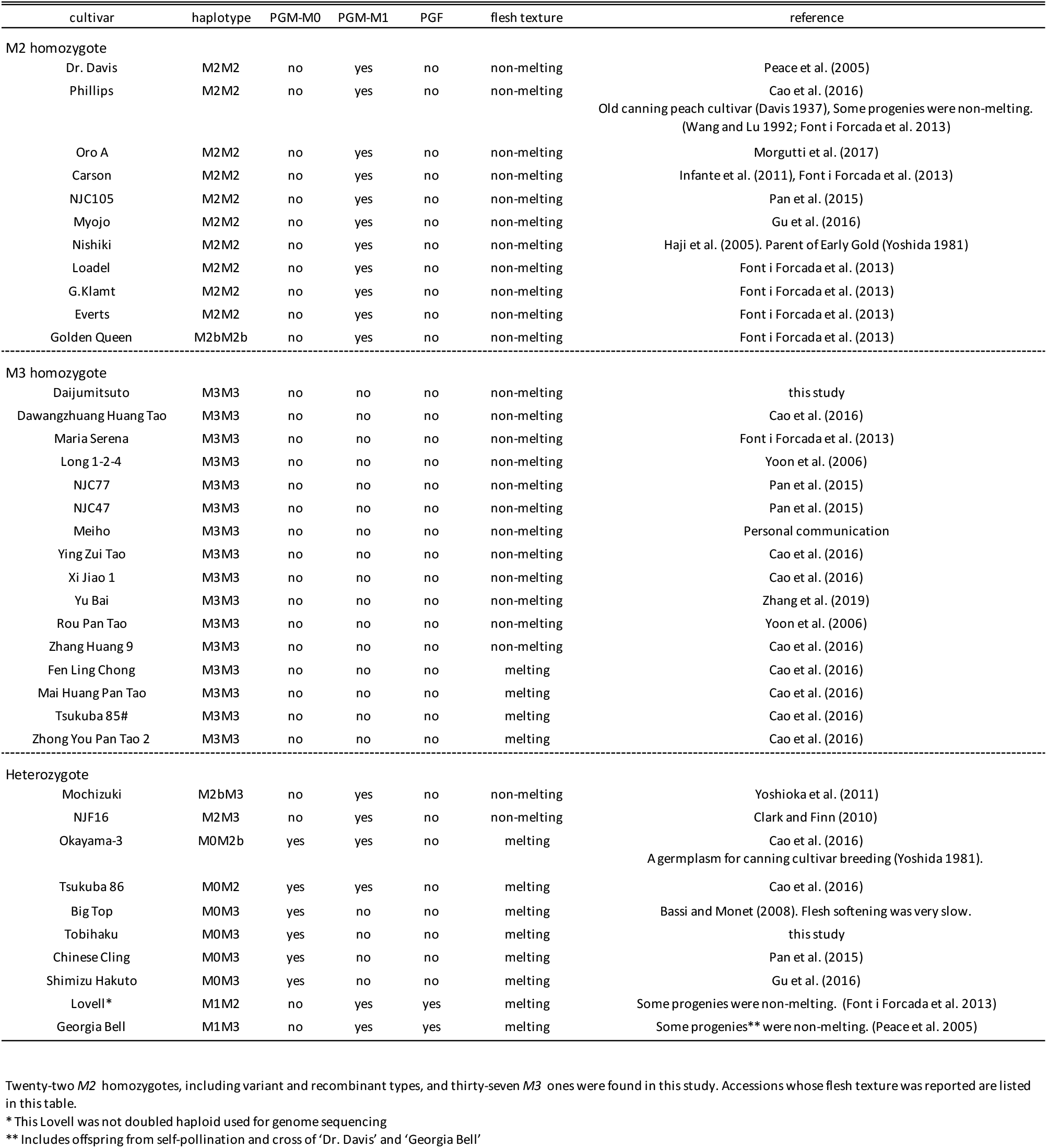
Flesh texture of *M^2^* or *M^3^* homozygotes and heterozygotes

| cultivar | haplotype | PGM-M0 | PGM-M1 | PGF | flesh texture | reference |
| --- | --- | --- | --- | --- | --- | --- |
| **M2 homozygote** |  |  |  |  |  |  |
| Dr. Davis | M2M2 | no | yes | no | non-melting | Peace et al. (2005) |
| Phillips | M2M2 | no | yes | no | non-melting | Cao et al. (2016) |
|  |  |  |  |  |  | Old canning peach cultivar (Davis 1937), Some progenies were non-melting.<br>(Wang and Lu 1992; Font i Forcada et al. 2013) |
| Oro A | M2M2 | no | yes | no | non-melting | Morgutti et al. (2017) |
| Carson | M2M2 | no | yes | no | non-melting | Infante et al. (2011), Font i Forcada et al. (2013) |
| NJC105 | M2M2 | no | yes | no | non-melting | Pan et al. (2015) |
| Myojo | M2M2 | no | yes | no | non-melting | Gu et al. (2016) |
| Nishiki | M2M2 | no | yes | no | non-melting | Haji et al. (2005). Parent of Early Gold (Yoshida 1981) |
| Loadel | M2M2 | no | yes | no | non-melting | Font i Forcada et al. (2013) |
| G.Klamt | M2M2 | no | yes | no | non-melting | Font i Forcada et al. (2013) |
| Everts | M2M2 | no | yes | no | non-melting | Font i Forcada et al. (2013) |
| Golden Queen | M2bM2b | no | yes | no | non-melting | Font i Forcada et al. (2013) |
| **M3 homozygote** |  |  |  |  |  |  |
| Daijimitsuto | M3M3 | no | no | no | non-melting | this study |
| Dawangzhuang Huang Tao | M3M3 | no | no | no | non-melting | Cao et al. (2016) |
| Maria Serena | M3M3 | no | no | no | non-melting | Font i Forcada et al. (2013) |
| Long 1-2-4 | M3M3 | no | no | no | non-melting | Yoon et al. (2006) |
| NJC77 | M3M3 | no | no | no | non-melting | Pan et al. (2015) |
| NJC47 | M3M3 | no | no | no | non-melting | Pan et al. (2015) |
| Meiho | M3M3 | no | no | no | non-melting | Personal communication |
| Ying Zui Tao | M3M3 | no | no | no | non-melting | Cao et al. (2016) |
| Xi Jiao 1 | M3M3 | no | no | no | non-melting | Cao et al. (2016) |
| Yu Bai | M3M3 | no | no | no | non-melting | Zhang et al. (2019) |
| Rou Pan Tao | M3M3 | no | no | no | non-melting | Yoon et al. (2006) |
| Zhang Huang 9 | M3M3 | no | no | no | non-melting | Cao et al. (2016) |
| Fen Ling Chong | M3M3 | no | no | no | melting | Cao et al. (2016) |
| Mai Huang Pan Tao | M3M3 | no | no | no | melting | Cao et al. (2016) |
| Tsukuba 85# | M3M3 | no | no | no | melting | Cao et al. (2016) |
| Zhong You Pan Tao 2 | M3M3 | no | no | no | melting | Cao et al. (2016) |
| **Heterozygote** |  |  |  |  |  |  |
| Mochizuki | M2bM3 | no | yes | no | non-melting | Yoshioka et al. (2011) |
| NJF16 | M2M3 | no | yes | no | non-melting | Clark and Finn (2010) |
| Okayama-3 | M0M2b | yes | yes | no | melting | Cao et al. (2016) |
|  |  |  |  |  |  | A germplasm for canning cultivar breeding (Yoshida 1981). |
| Tsukuba 86 | M0M2 | yes | yes | no | melting | Cao et al. (2016) |
| Big Top | M0M3 | yes | no | no | melting | Bassi and Monet (2008). Flesh softening was very slow. |
| Tobihaku | M0M3 | yes | no | no | melting | this study |
| Chinese Cling | M0M3 | yes | no | no | melting | Pan et al. (2015) |
| Shimizu Hakuto | M0M3 | yes | no | no | melting | Gu et al. (2016) |
| Lovell* | M1M2 | no | yes | yes | melting | Some progenies were non-melting. (Font i Forcada et al. 2013) |
| Georgia Bell | M1M3 | no | yes | yes | melting | Some progenies** were non-melting. (Peace et al. 2005) |

Twenty-two *M2* homozygotes, including variant and recombinant types, and thirty-seven *M3* ones were found in this study. Accessions whose flesh texture was reported are listed in this table.
\* This Lovell was not doubled haploid used for genome sequencing
\*\* Includes offspring from self-pollination and cross of 'Dr. Davis' and 'Georgia Bell'

### *M* Locus Structure in *Prunus* Species

The structure of *M* locus was identified from reference genome sequences of other *Prunus* species, including *P. mira, P. kansuensis*, almond (*P. dulcis*), apricot (*P. armeniaca*), Japanese apricot (*P. mume*), sweet cherry (*P. avium*), and Yoshino cherry (*P. yedoensis;* called ‘Sakura’ in Japan), and compared with the four main haplotypes of peach structurally identified in this study (Figs. 9 and S11). *PGM* and *PGF* were found in the Lauranne genome (PdLN) of almond, which belongs to subgenus *Amygdalus* together with peach, indicating that the PdLN *M* haplotype was similar to peach *M^1^* haplotype. As shown in *M^1^* of peach, *NADH* located downstream of *PGM* in PdLN was broken by an insertion sequence, although this insertion sequence was not similar to the M1 insertion. *F-box* and *NADH* were pseudogenes, probably due to multiple genome rearrangements occurring in almond Texas genome (PdTX). One *endoPG* was found in addition to *PG1* and *PG2*, and it was expected to be *PGF*, because *PGFs* in PdLN and PdTX shared amino acid substitutions specific to them and genomic sequence similarity was found not only in the gene region, but also in the intergenic region flanking them. A frameshift mutation was found in *PdLN-PGF*, indicating that *PdLN-PGF* did not function. The absence of this mutation in PdTX suggests that the mutation in *PdLN-PGF* occurred after the haplotype diverged. No tandem duplications of *PGM, PGF*, and M1 insertions were observed in *P. mira* or *P. kansuensis. NADH* appeared to be a fragmented structure in *P. kansuensis*, but because disrupted exons were located at different contigs in the draft genome, their actual relationship was unclear. *PGM, PGF*, and *NADH* were not present in *P. mira* genome.

**Figure 9.**
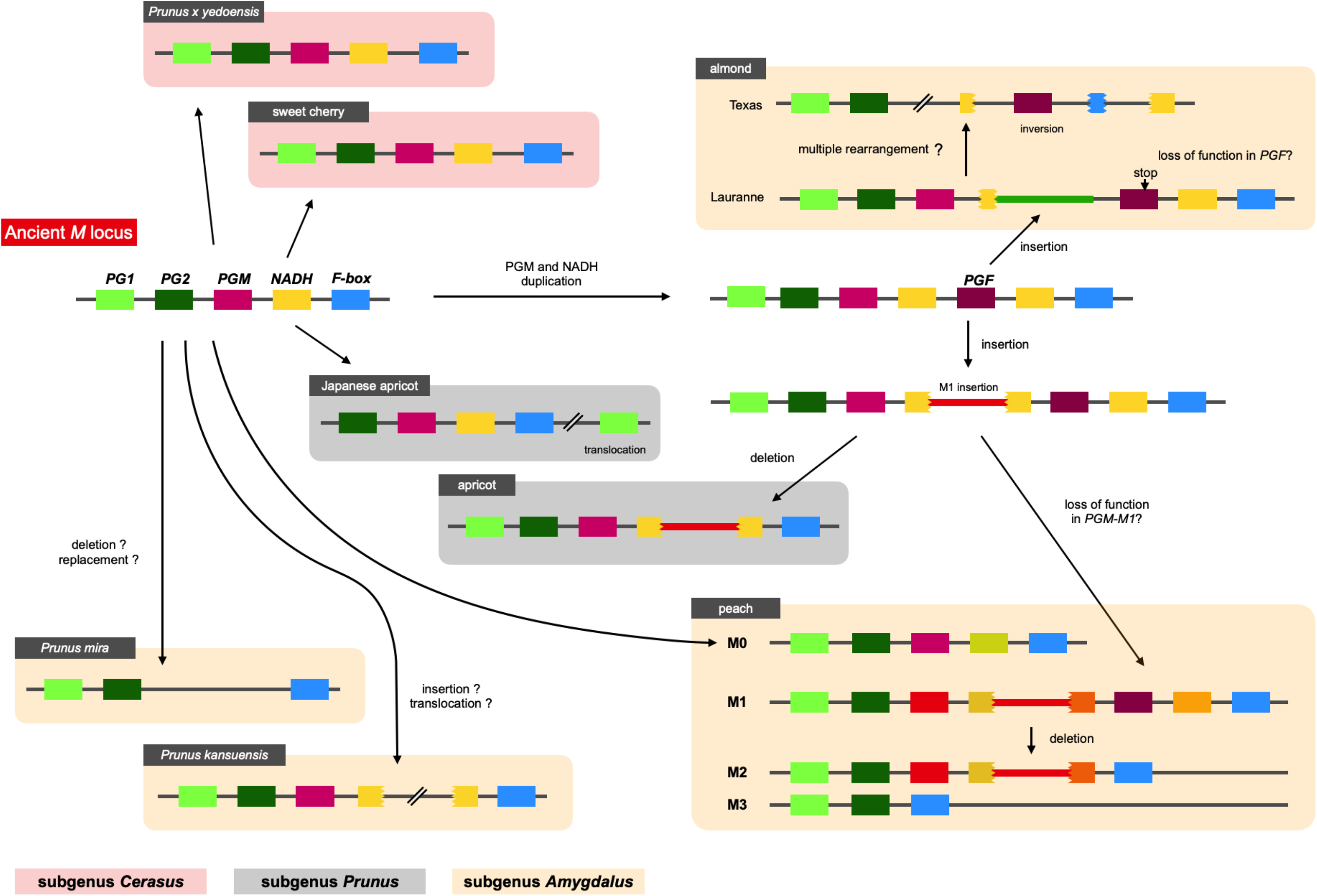
Diversification of *M* haplotypes in genus *Prunus*. We investigated *M* loci of *Prunus* species by Blastn analysis using their reference genomes. Five genes, *PG1, PG2, PGM, NADH*, and *F-box* genes, were shared among *Prunus* and speculated to be located at an ancient *M* haplotype. The structure of the ancient haplotype was conserved in peach M0, *P. kansuensis*, Japanese apricot, sweet cherry, and *P. yedoensis*. No *PGM* and *NADH* were observed in *P. mira. M^1^* type structure, which possessed *PGF* or M1 insertion, was found in peach *M^1^* and *M* haplotypes of almond and apricot. This haplotype was speculated to diverge from ancient haplotype by duplication of *PGM* (and *NADH*), followed by *M^1^* or almond insertion, which disrupted *NADH* gene located downstream of *PGM*.

In subgenus *Amygdalus*, the M1 insertion was found only in peach. However, a sequence similar to the M1 insertion was found in apricot of subgenus *Prunus*. The insertion also disrupted *NADH*, which was located downstream of *PGM*. These findings implied that the two insertions originated from the same event (haplotype), and that this insertion event occurred before the divergence of subgenera *Amygdalus* and *Prunus*. A structure similar to *M^0^* was found in Japanese apricot (*P. mume*) of subgenus *Prunus. PG2, PGM*, intact *NADH*, and *F-box* were clustered in *P. mume* haplotype, but *PG1* was translocated to the downstream region to *M* locus. Sweet cherry and Yoshino cherry belong to subgenus *Cerasus*, which is far from subgenus *Amygdalus*. Their *M* haplotype structure and gene composition were similar to those of peach *M^0^* haplotype.

## Discussion

### Different Postharvest Properties of TH and DJ

In this study, first, the postharvest properties of ultra-late maturing peach cultivars TH and DJ were investigated. Generally, peach is considered to be a climacteric fruit in which ripening-related phenomena, such as accelerated endogenous ethylene biosynthesis and fruit softening, were reported to be controlled by ethylene (Hayama et al., 2006; Liguori et al., 2004; Liu et al., 2018). TH exhibited the characteristics of normal MF peach fruit, including rapid fruit softening leading to MF texture associated with an increase in endogenous ethylene production. In contrast, DJ did not soften at all even though significant ethylene production was observed. It was considered that DJ fruit used in this study were harvested at optimum maturity, judging from their high sugar contents and low pH values at harvest. It seemed that DJ possessed the ability to produce ripening-related ethylene but lacked the ability to be softened in response to the ethylene. The lack of softening ability in response to ethylene in DJ was supported by the continuous propylene treatment. Propylene, instead of ethylene, treatment has been used to monitor endogenous ethylene production in parallel to other ripening-related changes. As much as 5,000 ppm of propylene was used for treatment in this study. According to previous reports, this concentration of propylene is equivalent to 50 ppm of ethylene and is sufficient to induce ethylene response in climacteric fruits (McMurchie et al., 1972; Burg and Burg, 1967). Indeed, in peach fruit treated with 500 ppm to 5,000 ppm of propylene, induction of autocatalytic ethylene production and dramatic fruit softening were reported (Liu et al., 2018; Yoshioka et al., 2010). In this study, DJ exhibited only a slight decrease in flesh firmness and maintained almost similar firmness to that at harvest even after seven days of continuous propylene treatment. As TH showed dramatic softening by the propylene treatment, it is suggested that the propylene treatment in this study is capable of inducing ethylene response in peach and that DJ has a non-softening characteristic in response to both endogenous and exogenous ethylene/propylene.

### Involvement of *M* locus in Determining Different Postharvest Properties of DJ and TH

The genetic background of DJ and TH and its relationships with other cultivars having long storability and/or shelf lives are unknown. One of the peach strains reported to have a long shelf life is SH peach (Haji et al., 2005). SH peaches, however, are characterized by the absence of ethylene production (Tatsuki et al., 2006). It is reported that ethylene sensing is normal in SH peaches and the fruit soften rapidly with exogenous ethylene or propylene treatment (Haji et al., 2003; Yoshioka et al., 2010), in contrast to DJ. A similar ripening characteristic observed in DJ has been reported in early harvest fruit of SR peaches (Brecht and Kader, 1984). In SR peaches harvested at an earlier date than the optimum harvest date, autocatalytic ethylene production was induced with or without propylene treatment whereas flesh firmness decreased quite slowly. However, DJ is phenotypically different from SR peaches in that it bears large fruit with red coloration, as shown in Figure 2, whereas SR fruit do not show normal ripening in terms of fruit size and coloration (Giné-Bordonaba et al., 2020). In agreement with these phenotypical differences between DJ and SH and/or SR peaches, the genomic sequences of DJ and TH did not exhibit any significant differences and/or mutations in the candidate causal genes for these specific strains, *YUCCA flavin mono-oxygenase* (Pan et al., 2015; Tatsuki et al., 2018) and *NAC transcription factor* (Eduardo et al., 2015; Meneses et al., 2016; Nuñez-Lillo et al., 2015) genes (data not shown).

On the other hand, significant differences between DJ and TH were found with regard to genomic sequences at *M* locus, which has been reported to control MF and NMF textures. *M* locus is composed of two tandem *endoPG* genes, *PGM* and *PGF* (Gu et al., 2016). Resequencing analysis revealed that DJ is a homozygote of the haplotype that lacks both *PGM*and *PGF*, designated as *M^3^* in this study, whereas TH is a heterozygote of*M^3^* and a structurally uncharacterized haplotype that possesses *PGM* (*PGM-M^0^*) but not *PGF*, designated as *M^0^* in this study (Figs. 4 and 5). Many reports have demonstrated that high level of enzymatic activity, protein accumulation, and gene expression of endoPG are observed only in MF, and supported the involvement of endoPG in the determination of the flesh texture (Reviewed in Bassi and Monet, 2008). Thus, it is considered that DJ is a member of NMF peaches. NMF peaches are known to be firm at maturity and to soften slowly during ripening without melting. Different from DJ, it was reported that softening progressed steadily during postharvest ripening in NMF peaches (Fishman et al., 1993; Yoshioka et al., 2011) and thus, the possible involvement of a number of factors in the non-softening property of DJ other than the lack of *endoPG* genes cannot be excluded. Nevertheless, it was indicated that the non-softening postharvest property of DJ is attributed to the lack of *endoPG* genes at *M* locus. The rapid softening property of TH is due to the presence of newly characterized *M^0^* haplotype (*PGM-M^0^* gene) in TH genome. This was further confirmed by the re-evaluation of *M*haplotype in relation to flesh textural phenotypes in 412 accessions, as described below.

### Re-evaluation of *M* Locus in Relation to Flesh Textural Phenotypes

In this study, we revealed that *M^0^* was not only a unique *M* haplotype as shown in TH, but also a widely spread haplotype in MF accessions, particularly popular peach cultivars grown in Japan, and was responsible for the MF phenotype in these accessions (Figs. 6 and 10B, Table S5). Based on the results obtained from the re-evaluation of *M* locus in 412 accessions in relation to flesh textural traits, we proposed the scenario in which four *M* alleles/haplotypes, *M^0^* to *M^3^*, were involved in the determination of flesh texture, with *M^0^* and *M^1^* dominantly controlling MF texture over *M^2^* and *M^3^*.

Various alleles/haplotypes have been proposed for *M* locus. The correspondence of *M* alleles/haplotypes in previous studies to those in this study is summarized in Tables S7 and S8. Peace et al. (2005) classified *endoPGs* at *M* locus into four alleles, F, f1, f, and null, on the basis of the results obtained from germplasm derived from peach cultivars GB and DD with f allele being hypothesized to be segregated via outcross from an unknown origin. Morgutti et al. (2017) assumed the same four haplotypes F, f1, f, and f_null_ by referring to Peace’s classification and further found two variations, PG^SH^ and PG^BT^, in f haplotype. The haplotype structures of H_1_, H_2_, and H_3_ reported by Gu et al. (2016) corresponded to those of F, f1, and f_null_ alleles, respectively, although the haplotype corresponding to f was not reported by Gu et al. (2016). In this study, we structurally identified four main haplotypes *M^0^* to *M^3^*. Judging from S49F substitution and indel at the third intron detected between *PGM-M^0^* and *PGM-M^1^, M^0^* appeared to be identical to f haplotype. However, the derivation of f (*M^0^*) haplotype assumed in this study was different from that in Morgutti et al. (2017), in which f (*M^0^*) was expected to be derived from F (*M^1^*) via fl (*M^2^*). We detected sequence diversifications and large structural differences between *M^0^* and *M^1^*/*M^2^* and assumed that *M^0^* was not derived from *M^1^*/*M^2^* directly. This assumption was supported by a structural comparison of *M* loci of other *Prunus* species (Fig. 9). A similar specific structure to peach *M^1^*, such as M1 insertion and/or tandem duplication of *endoPG*, was found in *M* haplotypes of almond and apricot. On the other hand, there was no insertion to disrupt *NADH* in peach *M^0^* or *M* haplotypes of the Japanese apricot, sweet cherry, and *P. yedoensis*. These findings suggested that the ancestral haplotypes of *M^0^* and *M^1^* diverged relatively early, before subgenus divergence, and evolved independently of each other. On the other hand, we could not find any SNP-level variation among *M^1^, M^2^*, and *M^3^* (Table S4), suggesting that the divergence of *M^1^* into *M^2^* and *M^3^* occurred relatively recently after *P. persica* speciation. It was also suggested that *M^2^* did not lead to *M^1^* but rather *M^2^* was derived from *M^1^*. This might be supported by the fact that the frequencies of *M^2^* and *M^3^* were much lower than that of *M^1^* at least in the accessions investigated in this study (Fig. 6). In this study, we showed 11 haplotypes in total (Fig. 7), but more haplotypes are expected to exist. We only examined reference genomes in *Prunus* species other than *P. persica*. Considering the divergence of *M* haplotype before speciation, it would not be surprising to find other species harboring haplotypes similar to both *M^0^* and *M^1^* haplotypes.

Gu et al. (2016) did not consider the presence of *PGM-M^0^* (f allele) and defined H_2_ as the sole haplotype harboring *PGM* but not *PGF* because they attempted to distinguish each haplotype on the basis of copy number of *endoPG* genes quantified by qPCR. Therefore, not only *M^2^* but also *M^0^* was genotyped as H_2_ in Gu et al. (2016). This misgenotyping of *M^0^* as H_2_ produced results that included incongruity between genotype and phenotype as *M^0^* (*PGM-M^0^*) and *M^2^* (*PGM-M^1^*) were likely to have different effects on flesh texture. For example, the genotyping in Gu et al. (2016) identified that SM and ‘Hakuho’ (HH), both of which are popular MF cultivars in Japan, were H_2_H_3_ and H_2_H_2_, respectively. Supposing *PGM-M^1^* on H_2_ is not functional, as assumed in this study, SM and HH should be NMF. Conversely, supposing H_2_ (*M^2^*) is a dominant haplotype that determines MF texture, as assumed by Gu et al. (2016), the H_2_H_2_ genotype shown in DD, ‘OroA’ (OA), and MZ cannot explain their NMF phenotypes.

Re-evaluation of *M* locus in association with MF/NMF phenotypes in this study revealed that *M^0^* and *M^1^* were likely to function dominantly over *M^2^* and *M^3^*. To our knowledge, *M^0^M^3^* was linked to MF, as shown in TH, SM, and BT. LL (*M^1^M^2^*) was also reported to exhibit MF phenotype (Font i Forcada et al., 2013), whereas NJF16 (*M^2^M^3^*) and MZ (*M^2b^M^3^*) had NMF phenotype (Clark and Finn, 2010; Yoshioka et al., 2011) (Table 1). Although *PGM-M^1^* was present in *M^2^*, its expression level seemed to be suppressed as reported in OA, whose genotype was determined as *M^2^M^2^* in this study (Morgutti et al., 2006). Thus, the low expression of *PGM-M^1^* was consistent with the feature of *M^2^*, namely, recessive against *M^1^* and *M^0^*, and comparable to *M^3^*. The hypothesis cannot be excluded that *M^2^* haplotypes in NMF accessions are specific haplotypes possessing additional mutation(s) that result in the disruption of *M^2^* function. The sequences of *PGM-M^1^* and its surrounding region on *M^2^* and *M^1^* haplotypes were identical with each other. As it was predicted that *M^2^* was generated from *M^1^* by deletion of the region including *PGF*, it seemed reasonable to consider that *PGM-M^1^* had lost its function before the emergence of *M^2^* haplotype and, thus *M^2^* haplotype in general was not functional. This prediction was supported by Morgutti et al. (2017), who reported that all accessions harboring flfl (*M^2^M^2^*) or flf_null_ (*M^2^M^3^*) exhibited NMF phenotype, as well as previous reports showing the existence of accessions exhibiting NMF phenotype but not completely lacking *endoPG* genes (Callahan et al., 2004; Lester et al., 1994, 1996; Morgutti et al., 2006; Peace et al., 2005).

It was hypothesized that two tandem *endoPG* genes at *M* locus, *PGM* and *PGF*, were responsible for peach flesh texture regulation (Gu et al., 2016). In this study, we suggested that *PGM-M^0^* and *PGF* in particular would affect flesh textural quality whereas *PGM-M^1^* would have no effect. Considering the lack of sequence diversification and the relatively recent divergence between *M^1^* and *M^2^*, it would not be possible that *PGM-M^1^* on *M^1^* retains its function. Although 11 haplotypes were characterized in this study, it might not be necessary to identify correctly all the haplotypes in order to estimate flesh textural quality in a breeding program. Only an analysis to confirm the presence of *PGM-M^0^* and *PGF* should be sufficient. This means we only need to test whether the PGM/F primer set (Figs. 8, S9, and S10B) amplifies *PGM-M^0^* or *PGF* fragments to estimate MF/NMF phenotypes in individual accessions and progenies.

*M* locus is strongly linked to freestone/clingstone (*F*) locus. It seems that *PGM-M^0^* is not responsible for the determination of *F* trait because most *M^0^M^0^* and *M^0^M^3^* accessions have clingstones. Gu et al. (2016) hypothesized that not *PGM* but only *PGF* on H_1_ (*M^1^*) haplotype is associated with the freestone phenotype. In this study, we re-evaluated the relationship between *M* genotypes and reported freestone/clingstone phenotypes. We found some accessions harboring *M^1^* haplotype but being reported to have not freestone but clingstone phenotype (data not shown). Further studies are required to unravel the role of *PGF* in the regulation of stone adhesion.

The classification of *M* haplotypes based on genomic structure and the re-evaluation of *M* locus in association with flesh melting traits in this study are expected to provide valuable information for studies on controlling fruit softening and textural quality. For example, BT is a well-known cultivar having slow softening behavior, but reaches MF texture at full maturity (Bassi and Monet, 2008; Ghiani et al., 2011). The *M* genotype of BT was found to be *M^0^M^3^* in this study, which was the same as that of SM, a famous Japanese MF cultivar that softens rapidly. Therefore, we suggest that the slow softening behavior of BT is controlled by loci other than *M* locus. Even with the re-evaluated genotypes in this study, there are few incongruities between *M* genotypes and MF/NMF phenotypes (Table S6). These incongruities may be due to different conditions and definitions for phenotyping flesh texture among experiments and/or specific mutation(s) that occurred in particular accessions, as well as the involvement of other loci. Further studies addressing the reason for these incongruities are expected to uncover mechanism(s) that determine flesh texture and fruit softening and to improve our understanding of factors related to long shelf life.

## Conclusion

We found that two ultra-late maturing cultivars, DJ and TH, showed different postharvest properties. DJ did not soften at all during ripening in spite of significant ethylene production, whereas TH showed rapid fruit softening leading to MF texture. Resequencing analyses of DJ and TH demonstrated that DJ was a homozygote of *M* haplotype designated as *M^3^* and lacked two tandem *endoPG* genes, *PGM* and *PGF*, at *M* locus. On the other hand, TH was a heterozygote of *M^3^* and a structurally unidentified haplotype designated as *M^0^* that consisted of only *PGM-M^0^* and was responsible for determining MF texture. Further classification of *M* haplotypes in 412 peach accessions revealed four main haplotypes: *M^0^; M^1^* consisting of *PGM-M^1^* and *PGF*; *M^2^* consisting of *PGM-M^1^* and *M^3^*; and *M^0^* was widely spread among MF accessions. We proposed the scenario that combinations of *M^0^* to *M^3^* determined flesh texture, and *M^0^* and *M^1^* dominantly controlled MF texture over *M^2^* and *M^3^*. These suggested the possibility that *PGM-M^0^* and *PGF* could confer MF phenotype and *PGM-M^1^* of *M^1^* and *M^2^* haplotypes may have lost its function. This scenario was supported by the evolution history of each *M* haplotype assumed from the structural features of *M* locus in *Prunus* species, in which the ancestral haplotypes of *M^0^* and *M^1^* diverged before subgenus divergence and evolved independently of each other, whereas *M^2^* and *M^3^* were assumed to be derived from *M^1^* in recent age by deletion mutations.

## Author Contributions

RN and TK equally contributed to this work. RN, TK, KU, and FF designed the study and drafted the manuscript. YF, DK, and MS performed sampling and phenotyping. RN, TK, YF, KA, and SY investigated postharvest ethylene production and flesh firmness. KU, RN, TK, and TA analyzed the genomic sequences of peach accessions and determined the genotypes of *M* locus in various peach accessions. All authors have contributed to manuscript revision and have read and approved the submitted version.

## Non-standard Abbreviations

Melting flesh locus, *M* locus; melting flesh, MF; non-melting flesh, NMF; stony-hard, SH; slow-ripening, SR; soluble solids content, SSC; endo-polygalacturonase, endo-PG; nicotinamide adenine dinucleotide dehydrogenase, NADH; SRA, sequence read archive; whole genome shotgun sequencing, WGS; single nucleotide polymorphism, SNP; insertion/deletion, indel

