## Supplementary material for "Postharvest properties of ultra-late maturing peach cultivars and their attributions to *Melting Flesh* (*M*) locus: Re-evaluation of *M* locus in association with flesh texture": Table S

Nakano et al. Figure S1  Soluble solids contents and juice pH in postharvest TH and DJ

Soluble Solid Contents

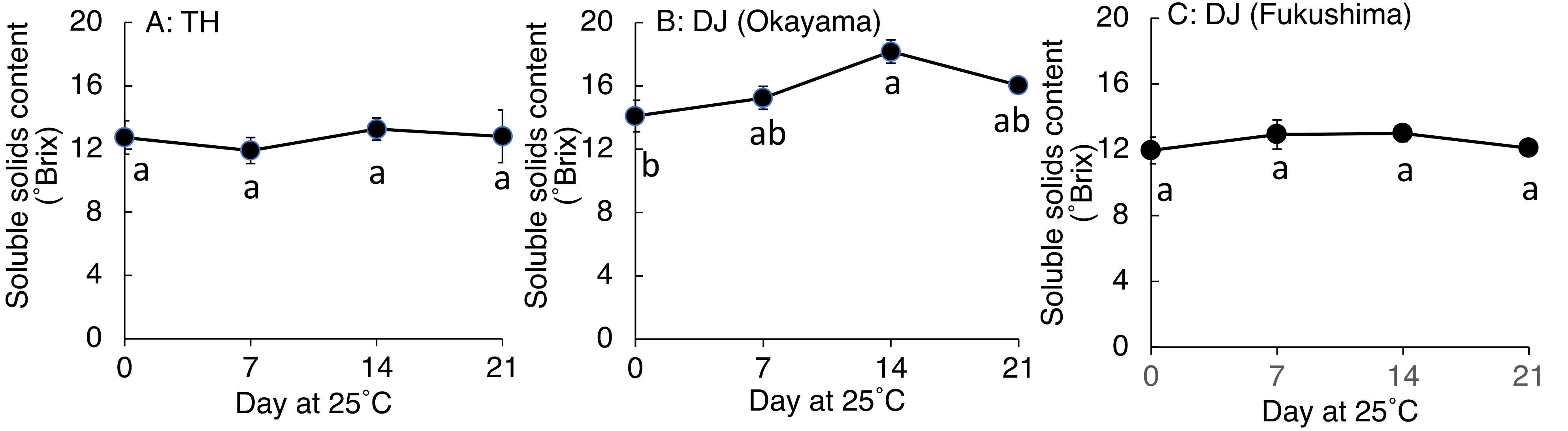


Juice pH

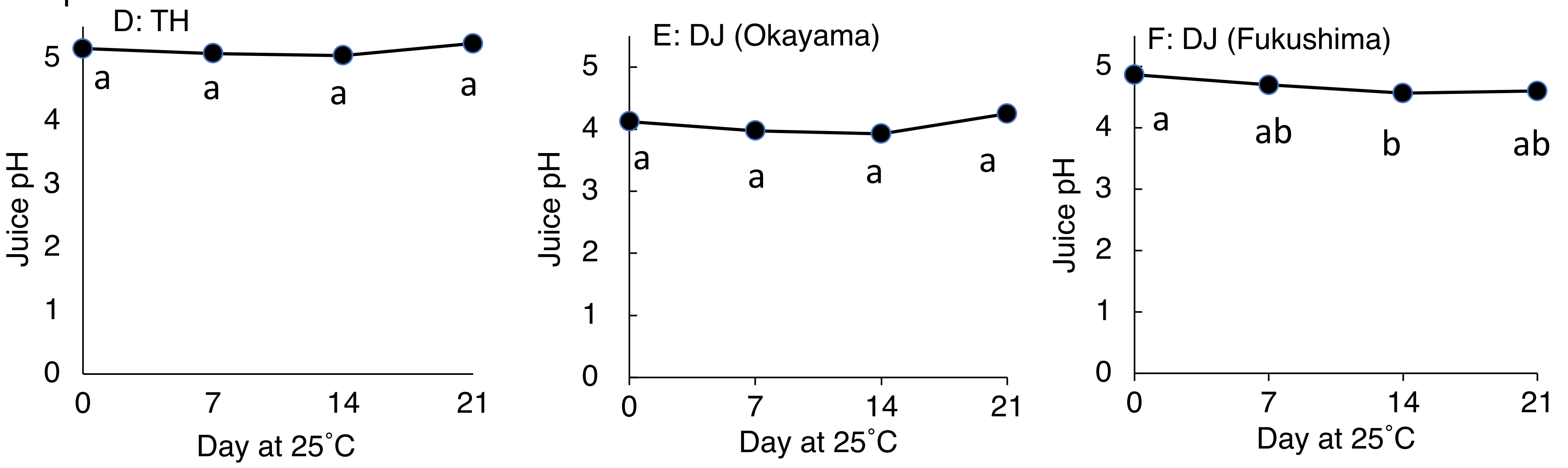

Nakano et al. Figure S2   Ethylene production in propylene treated TH and DJ

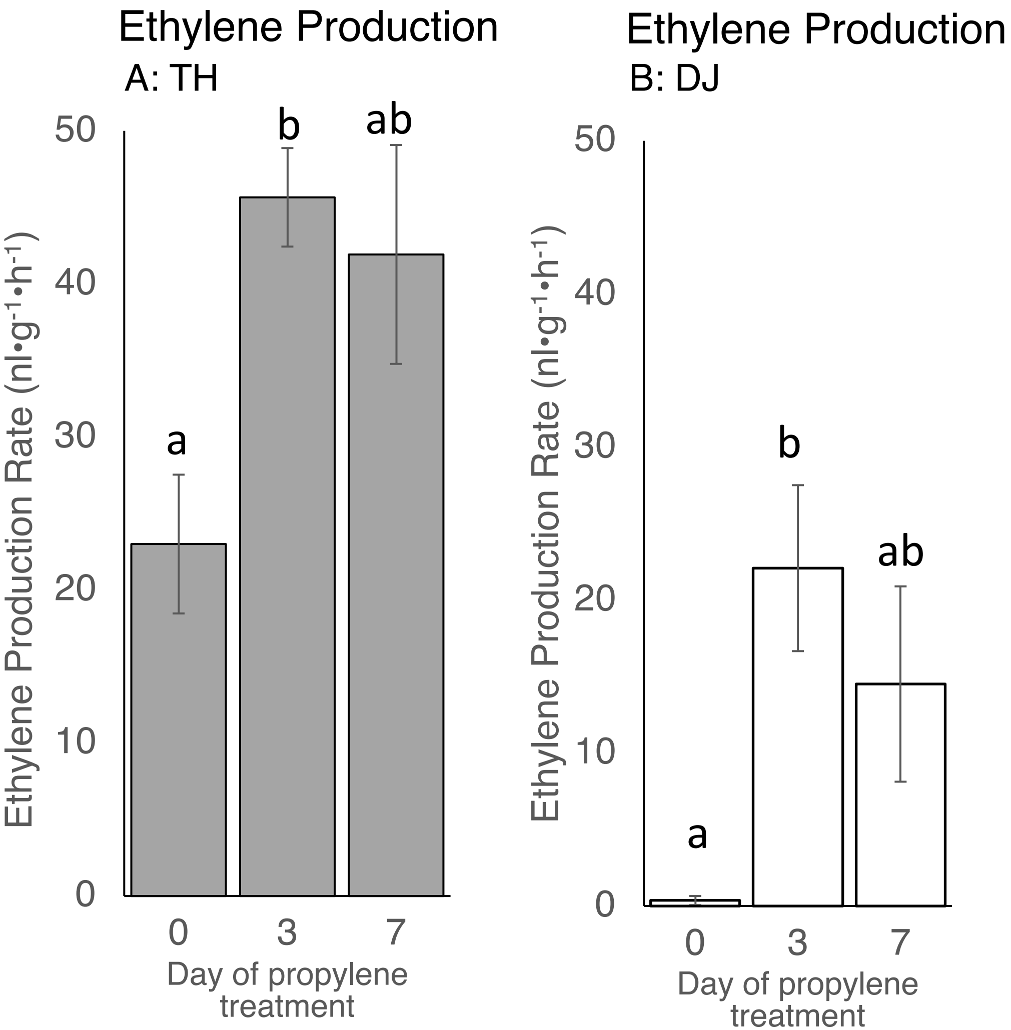

### NAKANO et al. Figure S3 Amino acid sequence comparison of *PGM-M*$^0$, *PGM-M*$^1$ and *PGF*.

```
                                        20                      40                      60                      80                     100
  PGM- M0  MANRRSLFSL SLIFVFMINS AIATPVTYNV ASLGAKADGK TDSTKAFLSA WAKACASMNP GVIYVPAGTF FLRDVVFSGP CKNNAITFRI AGTLVAPSDY 100
   PGM-M1  MANRRSLFSL SLIFVFMINS AIATPVTYNV ASLGAKADGK TDSTKAFLFA WAKACASMNP GVIYVPAGTF FLRDVVFSGP CKNNAITFRI AGTLVAPSDY 100
      PGF  MANRRSLFSL SLIFVFMINS AIATPVTYNV ASLGAKADGK TDSTKAFLSA WAKACASMNP GVIYVPAGTF FLRDVVFSGP CKNNAITFRI AGTLVAPSDY 100
 Pkan_PGM  MANRRSLFSL SLIFVFMINS AIATPVTYNV ASLGAKADGK TDSTKAFLSA WAKACASMNP GVIYVPAGTF FLRDVVFSGP CKNNAITFRI AGTLVAPSDY 100
 PdLN_PGM  MANRRSLFSL SLIFVFMINS AIATPVTYNV ASLGAKADGK TDSTKAFLSA WAKACASMNP GVIYVPAGTF FLRDVVFTGP CKNNAITFRI AGTLVAPSDY 100
 PdLN_PGF  MANRRSLLSL SLIFVFMINS AIATPVTYNV ASLGAKADGK TDSTKAFLSA WAKACASMNP GVIYVPAGTF FLRDVVFSGP CKNNAITFHI AGTLVAPSDY 100
 PdTX_PGF  MANRRSLLSL SLIFVFMINS AIATPVTYNV ASLGAKADGK TDSTKAFLSA WAKACASMNP GVIYVPAGTF FLRDVVFSGP CKNNAITFHI AGTLVAPSDY 100
 Pmum_PGM  MANRRSLFSL SLIFVFMINS AIATPVTYNV ASLGAKADGK TDSTKAFLSA WAKACASMNP GVIYVPAGTF FLRDVVFSGP CKNNAITFRI AGTLVAPSDY 100
 Parm_PGM  MANRRCLFSL SLIFVFMINS AIATPVTYNV ASLGAKADGK TDSTKAFLSA WAKACASMNP SVIYVPAGTF FLRDVVFSGP CKSNVITFRI AGTLVAPSDY 100
 Pavi_PGM  MANRRSLFSL SLVFVFMINS AIATPVTYNV ASLGAKADGK TDSTKAFLSA WAKACASMNP SVIYVPAGTF FLRDVVFSGP CKNNAITFRI AGTLVAPSDY 100
 Pyed_PGM  MANRRSLFSL ALVFVFMINS ATAAPVTYNV ASLGAKADGK TDSTKAFLSA WAKACASMNP GIIYVPAGTF FLRDVVFSGP CKNNAITFRI AGTLVAPSDY 100
                                       120                     140                     160                     180                     200
  PGM- M0  RVIGNAANWI FFHHVNGVTI SGGILDGQGT ALWACKASHG ESCPSGATTL GFSDSNNIVV SGLASLNSQM FHIVINDCQN VQMQGVRVSA SGNSPNTDGI 200
   PGM-M1  RVIGNAANWI FFHHVNGVTI SGGILDGQGT ALWACKASHG ESCPSGATTL GFSDSNNIVV SGLASLNSQM FHIVINDCQN VQMQGVRVSA SGNSPNTDGI 200
      PGF  RVIGNAANWI FFHHVNGVTI SGGILDGQGT ALWACKASHG ESCPSGATTL GFSDSNNIVV SGLASLNSQM FHIVINDCQN VQMQGVRVSA SGNSPNTDGI 200
 Pkan_PGM  RVIGNAANWI FFHHVNGVTI SGGILDGQGT ALWACKASHG ESCPSGATTL GFSDSNNIVV SGLASLNSQM FHIVINDCQN VQMQGVRVSA SGNSPNTDGI 200
 PdLN_PGM  RVIGNAANWI FFHHVNGVTI SGGILDGQGT ALWACKASHG ESCPSGATTL GFSDSNNIVV SGLASLNSQM FHIVINDCQN VQMQGVRVSA SGNSPNTDGI 200
 PdLN_PGF  RVIGNAANWI FFHHVNGVTI SGGILDGQGT ALWACKASHG ESCPSGATTL GFSDSNNIVV SGLASLNSQM FHIVINDCQN VQMQGVRVSA SGNSPNTDGI 200
 PdTX_PGF  RVIGNAANWI FFHHVNGVTI SGGILDGQGT ALWACKASHG ESCPSGATTL GFSDSNNIVV SGLASLNSQM FHIVINDCQN VQMQGVRVSA SGNSPNTDGI 200
 Pmum_PGM  RVIGNAANWI FFHHVNGVTI SGGILDGQGT ALWACKASNG KSCPSGATTL SFSDSNNIVV SGLASLNSQM FHIVINDCQN VQMQGVRVSA SGNSPNTDGI 200
 Parm_PGM  RVIGNAANWI FFHHVNGVTI SGGILDGQGT ALWACKASHG ESCPSGATTL SFSDSNNIVV SGLASLNSQM FHIVINDCQN VQMQGVRVSA SGNSPNTDGI 200
 Pavi_PGM  RVIGNAANWI FFHHVNGVTI SGGILDGQGT ALWACKASHG KSCPSGATTL GFSYSNNIVV SGLVSLNSQM FHIVINDCQN VQMQGVRVSA SGNSPNTDGI 200
 Pyed_PGM  RVIGNAAYWI FFHNVNGVTI SGGILNGQGT ALWACKASHG KSCPSGATTL GFSYSNNIVV SGLVSQNSQM FHIVINNCQN VQMQGVRVSA SGNSPNTDGI 200
                                       220                     240                     260                     280                     300
  PGM- M0  HVQMSSGVTI LNSKIATGDD CVSIGPGTSN LWIEGVACGP GHGISIGSLG KEQEEAGVQN VTVKTVTFTG TQNGLRIKSW GRPSTGFARN ILFQHATMVN 300
   PGM-M1  HVQMSSGVTI LNSKIATGDD CVSIGPGTSN LWIEGVACGP GHGISIGSLG KEQEEAGVQN VTVKTVTFTG TQNGLRIKSW GRPSTGFARN ILFQHATMVN 300
      PGF  HVQMSSGVTI LNSKIATGDD CVSIGPGTSN LWIEGVACGP GHGISIGSLG KEQEEAGVQN VTVKTVTFSG TQNGLRIKSW GRPSTGFARN ILFQHATMVN 300
 Pkan_PGM  HVQMSSGVTI LNSKIATGDD CVSIGPGTSN LWIEGVACGP GHGISIGSLG KEQEEAGVQN VTVKTVTFTG TQNGLRIKSW GRPSTGFARN ILFQHATMVN 300
 PdLN_PGM  HVQMSSGVTI LNSKIATGDD CVSIGPGTSN LWIEGVACGP GHGISIGSLG KEQEEAGVQN VTVKTVTFTG TQNGLRIKSW GRPSTGFARN ILFQHATMVN 300
 PdLN_PGF  HVQMSSGVTI LNSKIATGDD CVSIGSGTSN LWIEGVACGP GHGISIGKSR QGARRGRCTK CNS------- ---------- ---------- ---------- 263
 PdTX_PGF  HVQMSSGVTI LNSKIATGDD CVSIGPGTSN LWIEGVACGP GHGISIGSLG KEQEEAGVQN VTVKTVTFTG TQNGLRIKSW GRPSTGFARN ILFQHATMVN 300
 Pmum_PGM  HVQMSSGVTI LNSKIATGDD CVSIGPGTSN LWIESVACGP GHGISIGSLG KEQEEAGVQN VTVKTVTFTG TQNGLRIKSW GRPSTGFARN ILFQHATMVN 300
 Parm_PGM  HVQMSSGVTI LNSKIATGDD CISIGPGTSN LWIESVACGP GHGISIGSLG KEQEEAGVQN VTVKTVTFTG TQNGLRIKSW GRPSTGFARN ILFQHATMVN 300
 Pavi_PGM  HVQMSSGVTI LNSKIATGDD CVSIGPGTSN LWIESVACGP GHGISIGSLG KEQEEAGVQN VTVKTVTFTG TQNGLRIKSW GRASTGFARN ILFQHATMVN 300
 Pyed_PGM  HVQTSSGVTI LNSKIATGDD CVSIGPGTSN LWIESVACGP GHGISIGSLG KEQEEAGVQN VTVKTVTFTG TQNGLRIKSW GRPSTGFARN ILFQHATMVN 300
                                       320                     340                     360                     380
  PGM- M0  VENPIVIDQH YCPDNKGCPG QVSGVQISDV TYEDIHGTSA TEVAVKFDCS PKHPCSEIKL EDVKLTYKNQ AAESSCSHAD GTTEGVVQPT SCL 393
   PGM-M1  VENPIVIDQH YCPDNKGCPG QVSGVQISDV TYEDIHGTSA TEVAVKFDCS PKHPCSEIKL EDVKLTYKNQ AAESSCSHAD GTTEGVVQPT SCL 393
      PGF  VENPIVIDQH YCPDNKGCPG QVSGVQISDV TYEDIHGTSA TEVAVKFDCS PKHPCSEIKL EDVKLTYKNQ AAESSCSHAD GTTEGVVQPT SCL 393
 Pkan_PGM  VENPIVIDQH YCPNNKGCPG QVSGVQISDV TYEDIHGTSA TEVAVKFDCS PKHPCSEIKL EDVKLTYKNQ AAESSCSHAD GTTEGVVQPT SCL 393
 PdLN_PGM  VENPIVIDQH YCPDNKGCPG QVSGVQISDV TYEDIHGTSA TEVAVKFDCS PKYPCSEIKL EDVKLTYKNQ AAESSCSHAD GTTEGVVQPT SCL 393
 PdLN_PGF  ---------- ---------- ---------- ---------- ---------- ---------- ---------- ---------- ---------- --- 263
 PdTX_PGF  VENPIVIDQH YCPDNKGCPG QVSGVQISDV TYEDIHGTSA TEVAVKFDCS PKYPCSEIKL EDVKLTYKNQ AAESSCSHAD GTTEGVVQPT SCL 393
 Pmum_PGM  VENPIIIDQH YCPDNKGCPG QVSGVQISDV TYEDIHGTSA TEVAVKFDCS PEHPCSEIKL EDVKLTYKNQ AAESSCSHAD GTTEGVVQPT SCL 393
 Parm_PGM  VENPIIIDQH YCPDNKGCPG QVSGVQISDV TYEDIHGTSA TEVAVKFDCS PEHPCSEIKL EDVKLTYKNQ AAESSCSHAD GTTGGVVQPT SCL 393
 Pavi_PGM  VENPIVIDQH YCPDNKGCPG QVSGVQISDV TYEDIHGTSA TEVAVKFDCS PKHPCSEIKL KDVKLTYQNQ AAESSCSHAD GTTEGVVQPT SCL 393
 Pyed_PGM  VKNPIVIDQH YCPDNKGCPG QVSGIQISDV TYEDIHGTSA TEVAVKFDCS PKHPCSEIKL KDVKLTYQNK AAESSCSHAD GTTEGVVQPT SCL 393
```

# NAKANO et al.   Figure S5   Amino acid sequence comparison of NADH

```
                                                        exon1
        <--------------------------------------------------------------------------------------------------
NADH0   -----MAEVS NKQVILREYI TGFPKESDLY VSSTATIKLK LSEPPEHESS SKKLVLVKNL YLSCDPYQRL FMERIEGLSS QSTSSYTPGS PIYGYGVAKV 95
NADH1   ---------- ---------- ---------- ---------- ---------- ---------- ---------- ---------- -MKSALTKQN KTKKQHPIYG 19
NADH2   ---------- ---------- ---------- ---------- ---------- ---------- ---------- ---------- ---------- ---------- -
NADH3   -----MAEVS NKQVIPREYV TGFPKESDLY VNSTATIKLK LSEAPEHEGS SKKLVLVKNL YLSCDPYQRL FMERIEGLSS QSTSSYTPGS PIYGYGVAKV 95
NADH3/0 -----MAEVS NKQVIPREYV TGFPKESDLY VNSTATIKLK LSEAPEHEGS SKKLVLVKNL YLSCDPYQRL FMERIEGLSS QSTSSYTPGS PIYGYGVAKV 95
NADH3/1 -----MAEVS NKQVIPREYV TGFPKESDLY VNSTATIKLK LSEAPEHEGS SKKLVLVKNL YLSCDPYQRL FMERIEGLSS QSTSSYTPGS PIYGYGVAKV 95
AtNADH  MATSGNATVA NKQVILRDYV TGFPKESDL- IFTDSTIDLK IPE------- GSKTVLVKNL YLSCDPYMRI RMGKPDPGTA ALAPHYIPGE PIYGFSVSKV 92

                                exon2                                                        exon3
        ---><---------------------------------------------------> <----------------------------------------------
NADH0   LDSGHPDLKA GDLVWGTTYW EEYSRIPEPE GLIKIQHTDV PLSYYTGILG MPGLTSYVGF YEICSPKKGE HVFISAAAGA VGQLVGQFAK LMGCYVVGSA 195
NADH1   YGVGHPDLKA GDLVWGTTYW EEYSRIPEPE GLIKIQHTDV PLSYYTGILG MPGLTAYVGF YEICSPKKGE HVFISAAAGA VGQLVGQFAK LMGCYVVGSA 119
NADH2   ---------- ---------- ---------- ---------- ---------- -----MVKIQ ISGLQPWRFE LIYSAVVGVV EAPPKLGLLQ QDLSTQNQKQ 45
NADH3   LDSGHPDLKA GDLVWGTTYW EEYSRIPEPE GLIKIQHTDV PLSYYTGILG MPGLTAYVGF YEICSPKKGE HVFISAAAGA VGQLVGQFAK LMGCYVVGSA 195
NADH3/0 LDSGHPDLKA GDLVWGTTYW EEYSRIPEPE GLIKIQHTDV PLSYYTGILG MPGLTSYVGF YEICSPKKGE HVFISAAAGA VGQLVGQFAK LMGCYVVGSA 195
NADH3/1 LDSGHPDLKA GDLVWGTTYW EEYSRIPEPE GLIKIQHTDV PLSYYTGILG MPGLTAYVGF YEICSPKKGE HVFISAAAGA VGQLVGQFAK LMGCYVVGSA 195
AtNADH  IDSGHPDYKK GDLLWGLVGW GEYSLITPDF SHYKIQHTDV PLSYYTGLLG MPGMTAYAGF YEICSPKKGE TVFVSAASGA VGQLVGQFAK IMGCYVVGSA 192

                           exon4                                                     exon5
        ---><---------------------------------------> <-------------------------------------------------------------
NADH0   GS-------- ---------- ---------- --------ID IYFENVGGKT LDAVLLNMRV HG-RIAVCGM ISQYNLDQAE GVTNLMHLVY KRIRLHGFSV 258
NADH1   GSKEKVDLLK NELGFDEAFN YKEETDLNAA FKRYFPEGID IYFENVGGKT LDAVLLNMRV HG-RIAVCGM VSQYNLDQAE RVTNLMHLVY KRIRLHRFSV 218
NADH2   EQQTQVDLLK NELGFDEAFN YKEEADLNAA FKRYFPEGID IYFENVGGKT LDAVLLNMRV HG-RIAVCGM ISQYNLDQAE GVTNLMHLV- ---------- 133
NADH3   GSKEKHVQKE EEEGNTFEKE EEEASSKFEQ KELISLKPRG LRYPMKATAN DS-------- ---------- ---------- ---------- ---------- 247
NADH3/0 GS-------- ---------- ---------- --------ID IYFENVGGKT LDAVLLNMRV HG-RIAVCGM ISQYNLDQAE GVTNLMHLVY KRIRLHGFSV 258
NADH3/1 GSKEKVDLLK NELGFDEAFN YKEETDLNAA FKRYFPEGID IYFENVGGKT LDAVLLNMRV HG-RIAVCGM VSQYNLDQAE RVTNLMHLVY KRIRLHRFSV 294
AtNADH  GSNEKVDLLK NKFGFDDAFN YKAEPDLNAA LKRCFPEGID IYFENVGGKM LDAVLLNMKL HG-RIAVCGM ISQYNLEDQE GVHNLANVIY KRIRIKGFVV 291

                           exon5
        -------------------------------------------------------------->
NADH0   RDHYHLHPKF VEFMLPYIRQ GKIVYVEDIV EGLESGPRAL VGLFKGLNFG KQVVDVSA- 316
NADH1   RDHYHLHPKF VEFMLPYIRQ GKIVYVEDIV EGLESGPRAL VGLFKGLNFG KQVVDVSA- 276
NADH2   -DHYHIHPKF VEFMLPYIRQ GKIVYVEDIV EGLESGPRAL VGLFKGLNFG KQVVDVSA- 190
NADH3   ---------- ---------- ---------- ---------- ---------- --------- 247
NADH3/0 RDHYHLHPKF VEFMLPYIRQ GKIVYVEDIV EGLESGPRAL VGLFKGLNFG KQVVDVSA- 316
NADH3/1 RDHYHLHPKF VEFMLPYIRQ GKIVYVEDIV EGLESGPRAL VGLFKGLNFG KQVVDVSA- 352
AtNADH  SDYFDKHLKF LDFVLPYIRE GKITYVEDVV EGLENGPSAL LGLFHGKNVG KQLIAVARE 350
```

NAKANO et al.   Figure S6   Comparison of *NADH* gene structure among *Prunus* species.

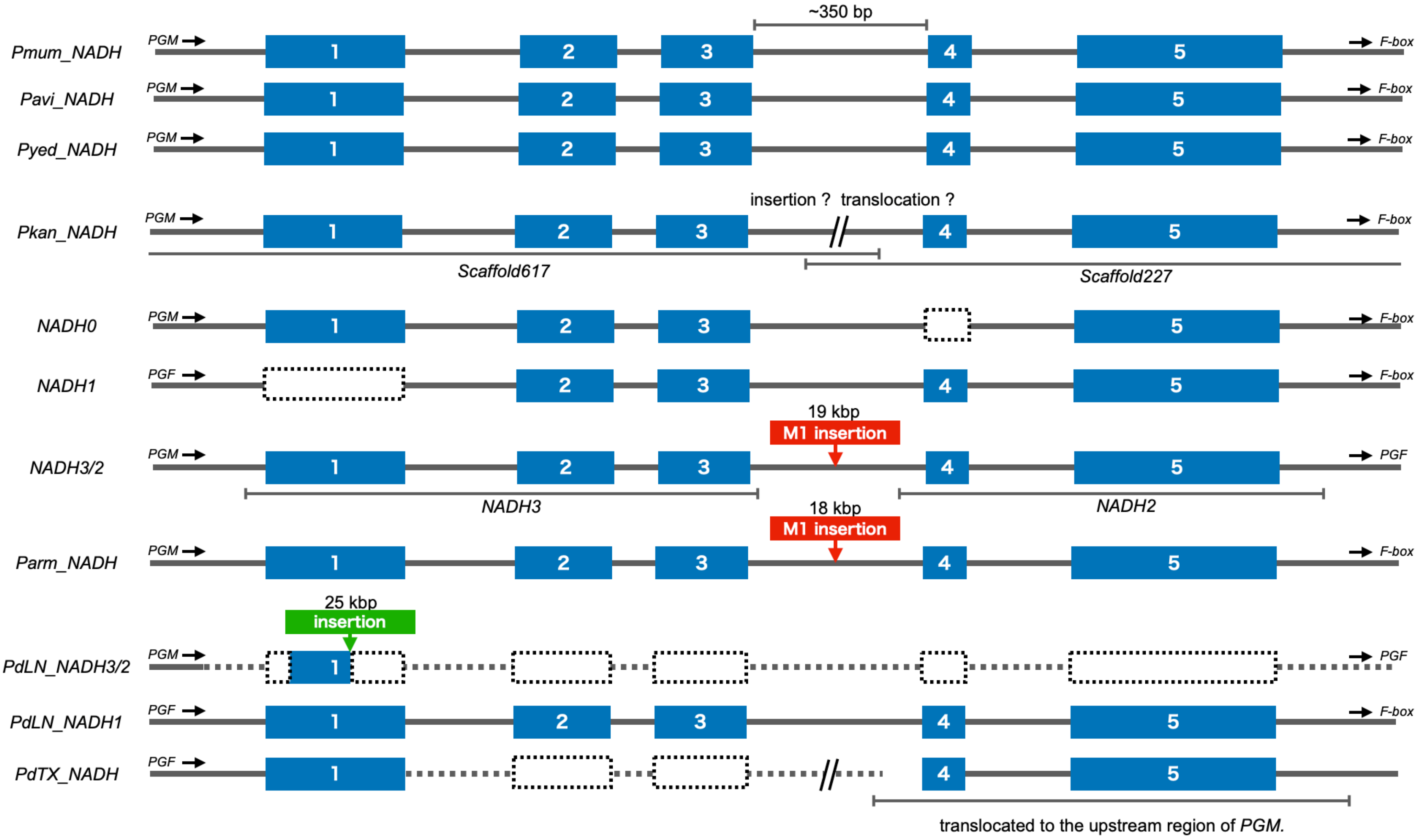

NAKANO et al.  Figure S9    MZ and O3 possessed $M^{2b}$ haplotype

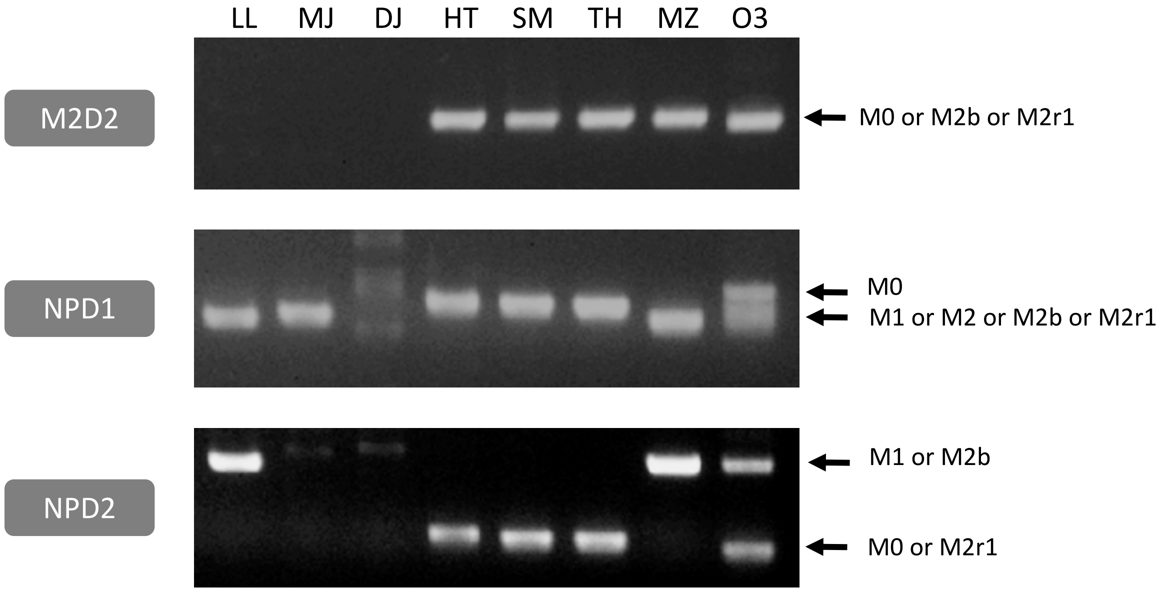

NAKANO et al. Figure S10 The peach cultivars bred in Japan shared an $M^0$ haplotype

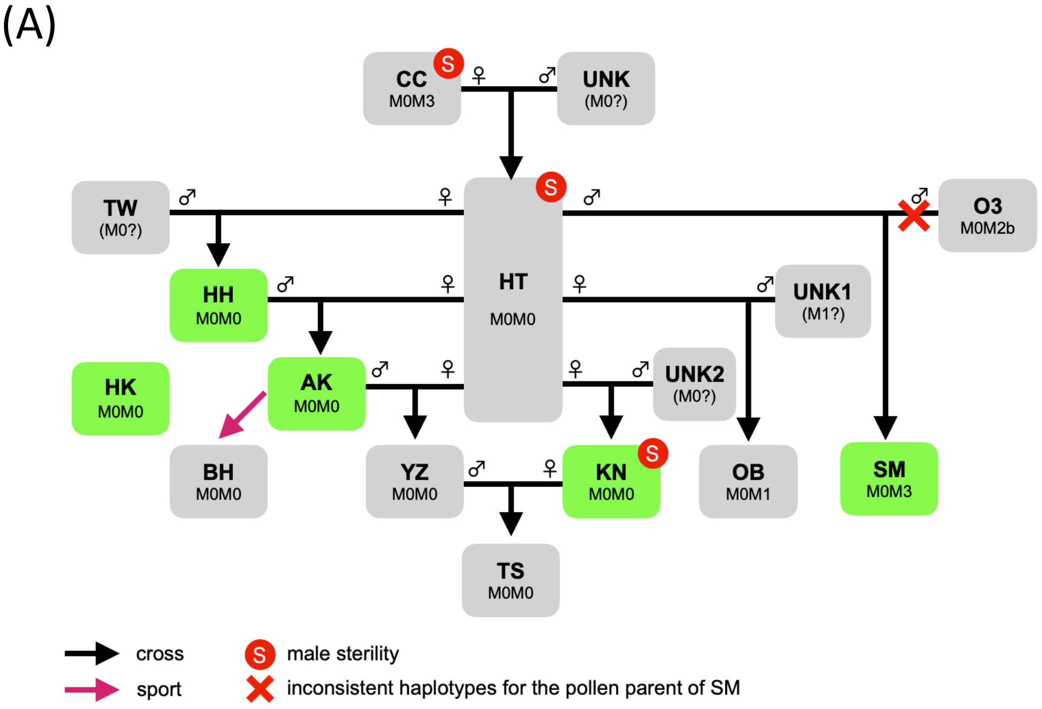


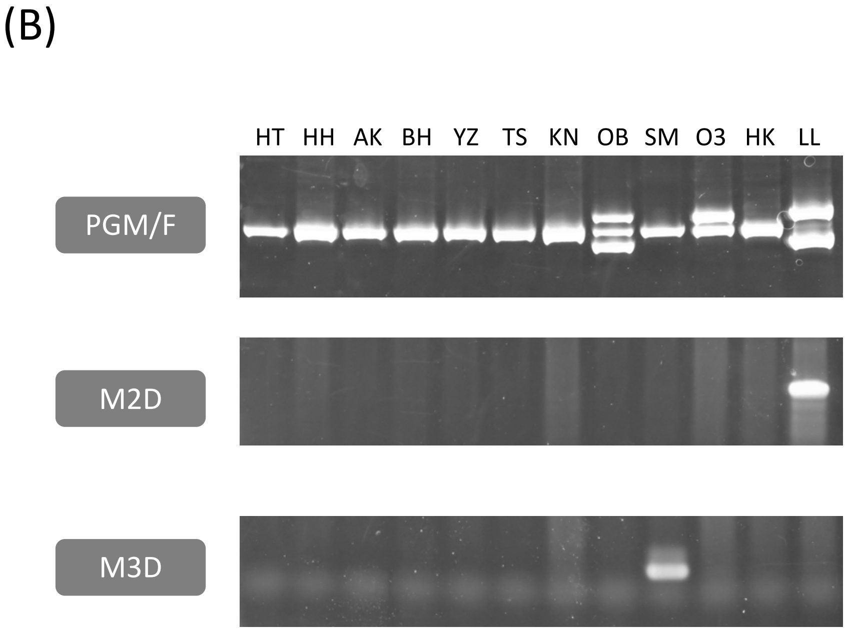

NAKANO et al. Figure S11 Structural comparison of *M* loci among *Prunus* species

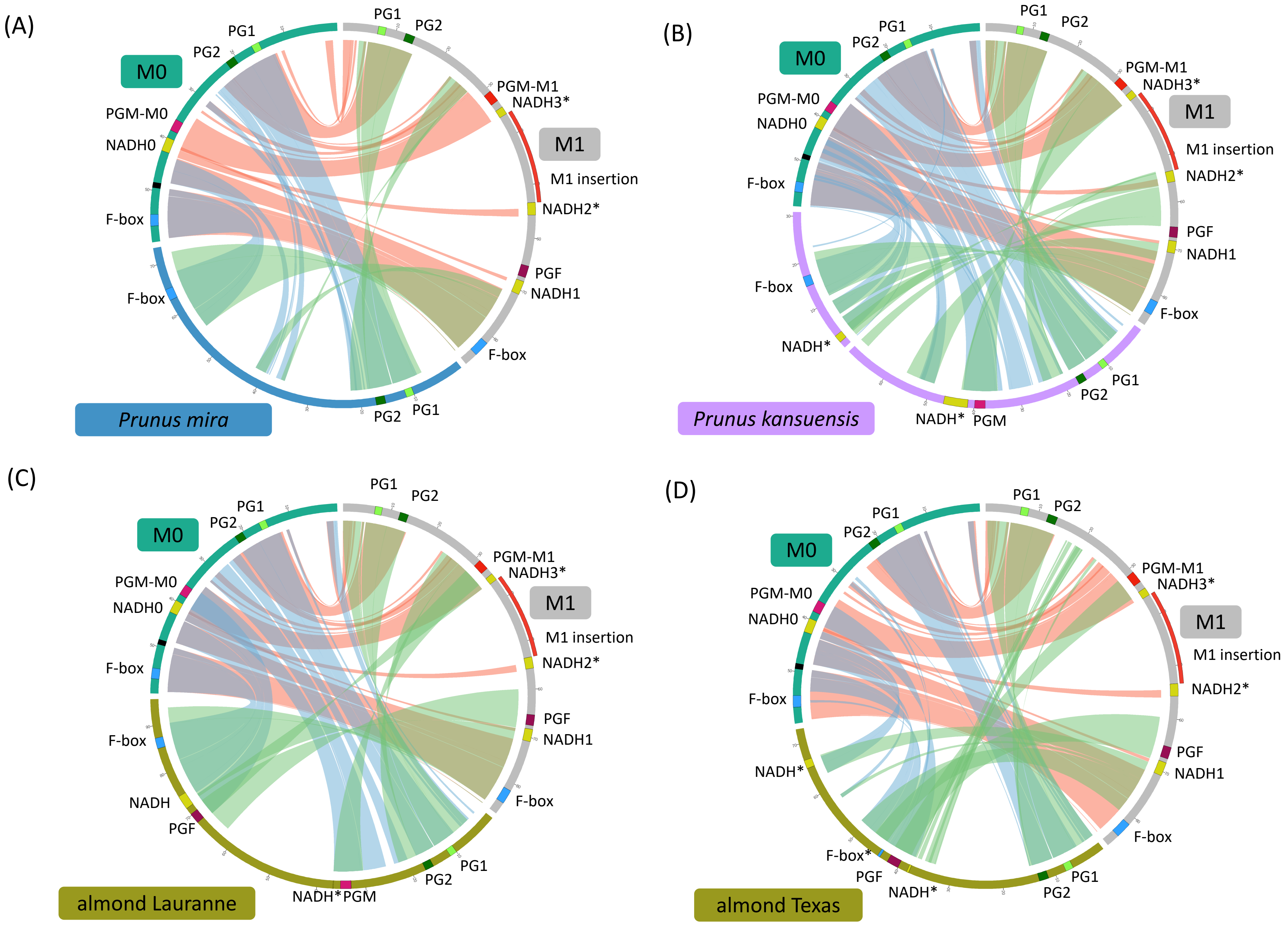

### NAKANO et al. Figure S11 Continued

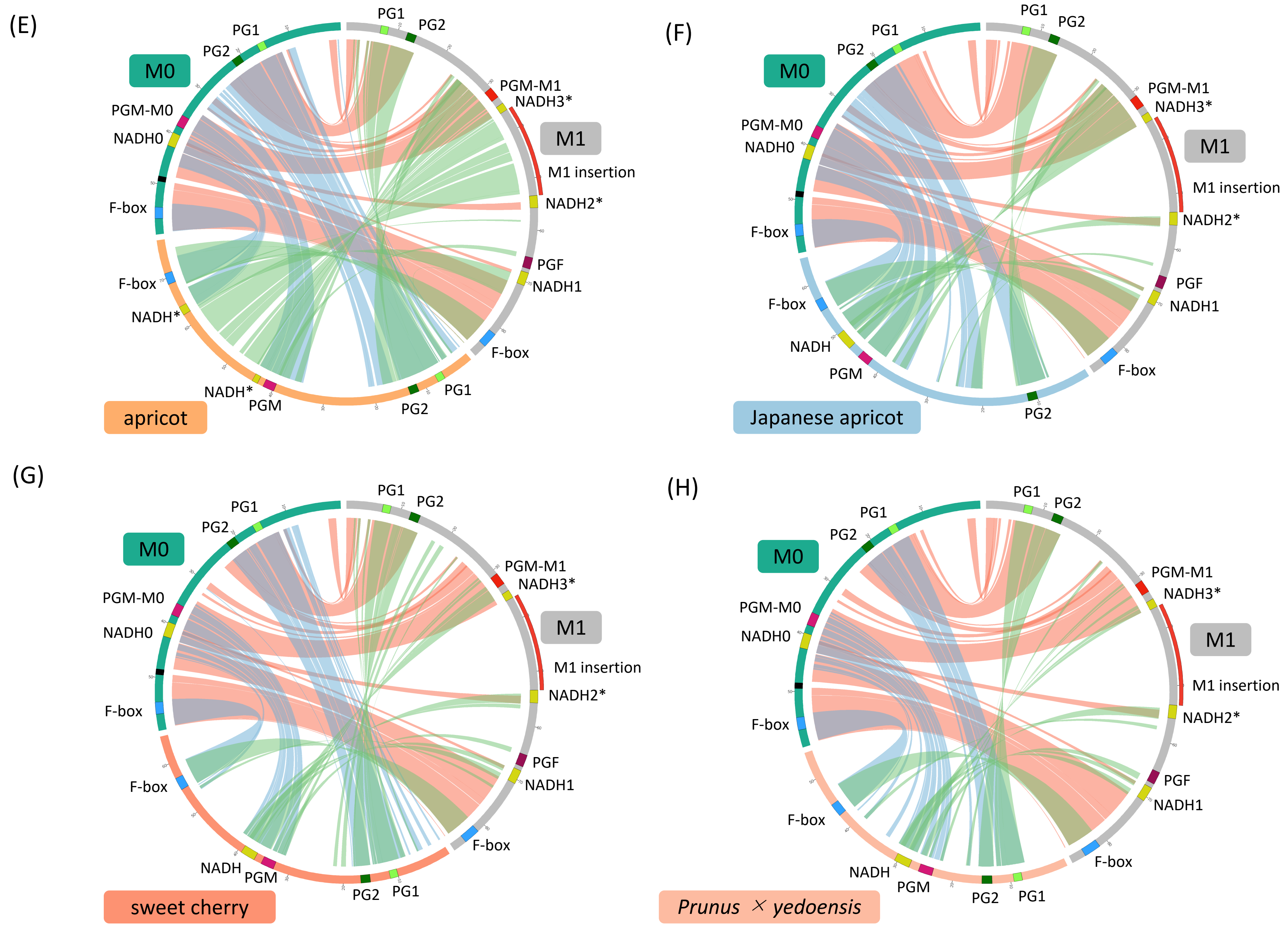
