## Supplementary material for "Postharvest properties of ultra-late maturing peach cultivars and their attributions to *Melting Flesh* (*M*) locus: Re-evaluation of *M* locus in association with flesh texture": Fig. S

Table S1. Climate conditions in peach production areas in Okayama and Fukushima Prefectures, Japan in 2018

| | Average Temperature (C) | | Cumulative Precipitation (mm) | | Average Humidity (%) | | Cumulative Hours of Solar Radiation (h) | |
|---|---|---|---|---|---|---|---|---|
| | Okayama | Fukushima | Okayama | Fukushima | Okayama | Fukushima | Okayama | Fukushima |
| Jan. | 3.5 | 1.4 | 37.0 | 72.5 | 71 | 70 | 155.6 | 142.1 |
| Feb. | 4.0 | 1.7 | 36.0 | 13.0 | 64 | 66 | 173.4 | 182.7 |
| Mar. | 10.1 | 7.8 | 147.5 | 93.5 | 67 | 59 | 231.0 | 220.1 |
| Apr. | 15.6 | 13.9 | 72.0 | 40.5 | 65 | 60 | 214.8 | 194.1 |
| May | 19.2 | 18.0 | 150.5 | 92.5 | 69 | 66 | 198.7 | 175.9 |
| Jun. | 22.9 | 21.6 | 140.0 | 26.0 | 74 | 70 | 169.4 | 182.9 |
| Jul. | 28.9 | 27.5 | 372.0 | 83.5 | 74 | 73 | 240.3 | 190.2 |
| Aug. | 29.3 | 26.3 | 37.0 | 105.5 | 67 | 74 | 259.6 | 159.2 |
| Sep. | 23.2 | 21.1 | 314.5 | 162.5 | 81 | 81 | 89.0 | 82.8 |
| Oct. | 18.2 | 16.2 | 40.0 | 69.5 | 71 | 76 | 198.8 | 131.6 |
| Nov. | 12.5 | 10.7 | 2.5 | 7.5 | 73 | 73 | 169.2 | 145.0 |
| Dec. | 7.7 | 4.4 | 61.0 | 61.5 | 76 | 74 | 129.4 | 109.1 |

Table S2. Relationships between flesh penetration force measured by the system used in this study and other fruit maturity indexes.

| Developmental stage | Flesh penetration force (N/mm2) | Epicarp color | Ethylene production | Characteristics |
|---|---|---|---|---|
| Unripe | >1.5 | Green | No ethylene detected | Immature fruit before onset of harvest |
| Physiologically ripe | 0.6-1.0 | Light green/yellow/red | Little ethylene produced | Commercial harvest maturity, slightly firm/crispy flesh |
| Fully ripe | <0.25 | Yellow/red | Much ethylene produced | Ready to eat, soft/melting flesh, easily damaged |

Table S4. Comparison of SNP number in regions from *PG1* to *PG2*

| cultivar | genotype | *PG1* | intergenic region 1 | intergenic region 2 | *PG2* |
|---|---|---|---|---|---|
| Daijyumitsuto | *M3M3* | 0 | 0 | NA | NA |
| Tobihaku | *M0M3* | 13(13) | 25(25) | 13(1) | 15(0) |
| Big Top | *M0M3* | 13(12) | 17(17) | 20(0) | 22(0) |
| Benihakuto | *M0M0* | 18(2) | 36(1) | 17(0) | 25(0) |
| Lovell | *M1M1* | 0 | 0 | 0 | 0 |
| Dr. Davis | *M2M2* | 0 | 0 | 0 | 0 |

Intergenic region 1 was from end of *PG1* to 19,026,185 bp (outside H3 deletion region).

Intergenic region 2 was from 19,026,186 bp (inside H3 deletion region) to start of *PG2* .

Values in parentheses are those of heterozygous SNPs.

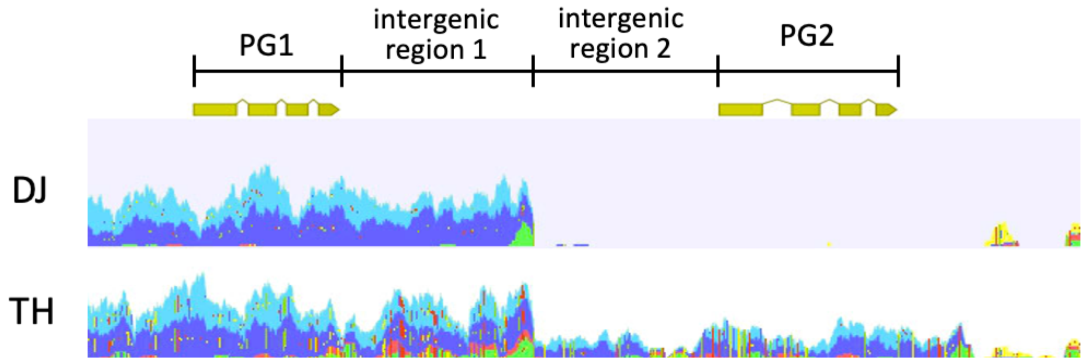

Table S6. Differences in flesh texture predicted by genotype and reported phenotype

| cultivar | genotype | phenotype predicted by genotype | reported phenotype | reference |
|---|---|---|---|---|
| Hakuho | *M0M0* | melting | melting* | Pan et al. (2015) |
| Nagasawa Hakuho | *M0M0* | melting | melting* | Gu et al. (2016) |
| Hakuho | *M0M0* | melting | non-melting | Yoon et al. (2006) |
| Nagasawa Hakuho | *M0M0* | melting | non-melting | Yoon et al. (2006) |
| Diao Zhi Bai | *M0M1* | melting | Non-melting | Yoon et al. (2006) |
| Shou Bai | *M1r2M1r2* | melting | Non-melting | Yoon et al. (2006) |
| Wu Yue Xian Bian Gan | *M0M0* | melting | non-melting | Yoon et al. (2006) |
| Fen Ling Chong | *M3M3* | non-melting | Melting | Cao et al. (2016) |
| Mai Huang Pan Tao | *M3M3* | non-melting | Melting | Cao et al. (2016) |
| Tsukuba 85 | *M3M3* | non-melting | Melting | Cao et al. (2016) |
| Zhong You Pan Tao 2 | *M3M3* | non-melting | Melting | Cao et al. (2016) |
| Zhong You Pan Tao 4 | *M0M3* | melting | non-melting | Cao et al. (2016) |
| Early Gold | *M1M1* | melting | non-melting | Yoshida (1981) |

* The reported phenotype matched the predicted one in this study.
The 11 accessions showed differences between predicted and reported phenotypes. All except 'Early Gold' (EG) were reported by Yoon et al. (2006) and Cao et al. (2016).
EG was a canning peach and its parent was 'Nishiki' (NK). EG should possess at least one *M2* haplotype because NK was an *M2* homozygote, but resequencing analysis showed that EG was M1M1.

Table S7. Correlation of *PGM/F* genes in this study with those in previous reports

| this study | Gu et al. (2016) | Morgutti et al. (2017) | Peace et al. (2005) |
|---|---|---|---|
| *PGM-M0* | not found | $PG^{SH}$, $PG^{BT}$ | f |
| *PGM-M1* | PGM | $PG^{m}$ | f1, F(a) |
| *PGF* | PGF | *PG_M* | F(b) |
| null | null | null | n(ull) |

$PG^{BT}$ in Morgutti et al. (2017) was *Bst* XI-sensitive (Fig. S6).

Table S8. Correlation of haplotypes in this study with those in previous reports

| haplotype | | | number of genes | | | | | | |
|---|---|---|---|---|---|---|---|---|---|
| this study | Gu et al. (2016) | Morgutti et al. (2017) | *PG1* | *PG2* | *PGM-M0* | *PGM-M1* | *PGF* | *NADH* | *F-box* |
| *M0* | not found* | f | 1 | 1 | 1 | 0 | 0 | 1 | 1 |
| *M1* | H1 | F | 1 | 1 | 0 | 1 | 1 | 3 | 1 |
| *M2* | H2 | f1 | 1 | 1 | 0 | 1 | 0 | 2 | 1 |
| *M3* | H3 | $f_{null}$ | 1 | 0 | 0 | 0 | 0 | 0 | 1 |

**M0* was misidentified as *H2* haplotype in Gu et al. (2016).

f haplotype structure in Morgutti et al. (2017) was postulated from *M1* haplotype. *PG* gene composition was corrected but the genome structure was not the same as *M0* haplotype in this study.

*PGM-M0* and *PGM -M1* were allelic.
